# Heterogeneous Cardiac- and Neural Crest-Derived Aortic Smooth Muscle Cells have Similar Transcriptional Changes after TGFβ Signaling Disruption

**DOI:** 10.1101/2024.04.28.591539

**Authors:** Pengwei Ren, Bo Jiang, Abdulrahman Hassab, Guangxin Li, Wei Li, Roland Assi, George Tellides

## Abstract

Smooth muscle cells (SMCs) of cardiac and neural crest origin contribute to the developing proximal aorta and are linked to disease propensity in adults. We analyzed single-cell transcriptomes of SMCs from mature thoracic aortas in mice to determine basal states and changes after disrupting transforming growth factor-β (TGFβ) signaling necessary for aortic homeostasis. A minority of Myh11 lineage-marked SMCs differentially expressed genes suggestive of embryological origin. Additional analyses in Nkx2-5 and Wnt1 lineage-marked SMCs derived from cardiac and neural crest progenitors, respectively, showed both lineages contributed to a major common cluster and each lineage to a minor distinct cluster. Common cluster SMCs extended from root to arch, cardiac subset cluster SMCs from root to mid-ascending, while neural crest subset cluster SMCs were restricted to the arch. The neural crest subset cluster had greater expression of a subgroup of TGFβ-dependent genes suggesting specific responsiveness or skewed extracellular matrix synthesis. Nonetheless, deletion of TGFβ receptors in SMCs resulted in similar transcriptional changes among all clusters, primarily decreased extracellular matrix molecules and modulators of TGFβ signaling. Many embryological markers of murine aortic SMCs were not confirmed in adult human aortas. We conclude: (i) there are multiple subtypes of cardiac- and neural crest-derived SMCs with shared or distinctive transcriptional profiles, (ii) neural crest subset SMCs with increased expression of certain TGFβ-inducible genes are not spatially linked to the aortic root predisposed to aneurysms from aberrant TGFβ signaling, and (iii) loss of TGFβ responses after receptor deletion is uniform among SMCs of different embryological origins.

## Introduction

Smooth muscle cells (SMCs), the major and often exclusive cell type of the thick medial layer of arteries, have diverse embryological origins depending on somatic location. Accordingly, the aorta which traverses the thorax and abdomen, consisting of root, ascending, arch, descending thoracic, and abdominal segments, contains SMCs from multiple sources (1). Studies in chimeric quail-chick embryos demonstrated that SMCs of arteries which derive from the pharyngeal arches (analogous to mammalian aortic arch and certain of its branches) are entirely of neural crest origin, those in a transition zone between the bulbus arteriosus and aortic trunk arising from the heart (analogous to mammalian aortic root and ascending aorta) are partly of neural crest and partly of mesodermal origin, and those of the dorsal aorta (analogous to mammalian descending aorta) are entirely of mesodermal origin (2). Additional investigations in quail-chick chimeras showed that mesodermal SMCs at the base of the aorta derive from the secondary heart field that contributes myocardium to the outflow tract (3). Genetic lineage tracing in mice confirmed a convoluted SMC distribution in which the aortic root and outer layers of the ascending aorta derive from second heart field progenitors, whereas the aortic arch and inner layers of the ascending aorta derive from cardiac neural crest progenitors (4,5). Ablation of secondary heart field or neural crest cells in avian models lead to conotruncal defects demonstrating non-redundant functions in great artery development (6,7).

In addition to associations with congenital anomalies, SMCs of different embryological origins are frequently invoked to explain disease focality in mature thoracic aortas of humans. Atherosclerosis preferentially afflicts the aortic arch and descending aorta, while aneurysms are more common in the aortic root and ascending aorta. In particular, Marfan and Loeys-Dietz syndromes due to heterozygous mutations in genes encoding fibrillin-1 and the transforming growth factor-β (TGFβ) signaling cascade, respectively, predominantly manifest as aortic root aneurysms (8). Mechanisms of aneurysm formation and predilection for specific segments, however, are incompletely understood and controversial. A leading hypothesis for both Marfan and Loeys-Dietz syndromes is pathogenic TGFβ overactivity (9), despite more severe disease after TGFβ inhibition in experimental models of the former (10–12) and inactivating mutations in TGFβ receptors in the latter (13). Lineage-specific events are a possible explanation for local differences in TGFβ activity. Indeed, cultured SMCs from chick embryo aortic arch show greater TGFβ transcriptional activity by luciferase reporter assay than from the abdominal aorta (14). In a murine model of Loeys-Dietz syndrome with a heterozygous inactivating mutation of *Tgfbr1*, paracrine interactions between signaling-deficient, cardiac-derived SMCs and signaling-sufficient, neural crest-derived SMCs result in aneurysms restricted to the aortic root (15). However, mechanisms that spare TGFβ signaling in neural crest-derived SMCs are not elucidated nor reasons for an absence of aneurysms in the ascending aorta where interactions of SMCs from different embryological origins are more widespread than in the aortic root.

We, therefore, sought to characterize SMCs of the adult thoracic aorta at single-cell level to determine basal molecular profiles of cardiac- and neural crest-derived SMCs and transcriptional responses after categorical disruption of TGFβ signaling.

## Results

### Minor Clusters of SMCs are Distinguished by Cardiac- and Neural Crest-Related Genes

To identify markers suggestive of embryological origin, single-cell RNA sequencing (scRNA-seq) was performed in SMCs isolated from root, ascending, and arch aortic segments of 12-week-old mice. The aortas were digested at 4 °C to minimize postmortem cellular stress and transcriptional artifact (16). An inducible Myh11-CreER;mT/mG dual reporter enabled selection of Myh11 lineage-marked SMCs expressing green fluorescent protein (GFP) and exclusion of other cell types expressing red fluorescent protein (RFP). Data from poor-quality SMCs were filtered using several RNA metrics (Supplemental Figure 1A-C). Unsupervised clustering of 5,238 cells from 2 replicates revealed 3 clusters of SMCs (Figure 1A,B). Differentially expressed genes were scrutinized for cardiac or neural crest associations (Supplemental Figure 1D). A minor cluster (#2) expressed several homeobox transcription factors, including *Prxx2*, *Dlx5*, *Dlx6*, and *Msx2*, related to neural crest development (17) (Figure 1C). Another minor cluster (#1) differentially expressed *Des* encoding desmin the major intermediate filament of cardiac but not aortic smooth muscle, *Tnnt2* encoding a cardiac and skeletal muscle-restricted component of the troponin complex, and a long noncoding RNA, *Hand2os1* and a homeobox transcription factor, *Meis2* described to regulate cardiac development (18–21) (Figure 1D). The major cluster (#0), however, was not distinguished by cardiac- or neural crest-related markers, while all 3 clusters expressed SMC-restricted genes, e.g., *Myh11* and *Itga8*, but not typical cardiomyocyte genes, e.g., *Myh7* and *Tnni3* (Figure 1E). These findings indicate that although a minority of root, ascending, and arch SMCs differentially express transcripts suggestive of embryological origins, most cannot be distinguished as cardiac- or neural crest-derived without further lineage tracing.

**Figure 1:**
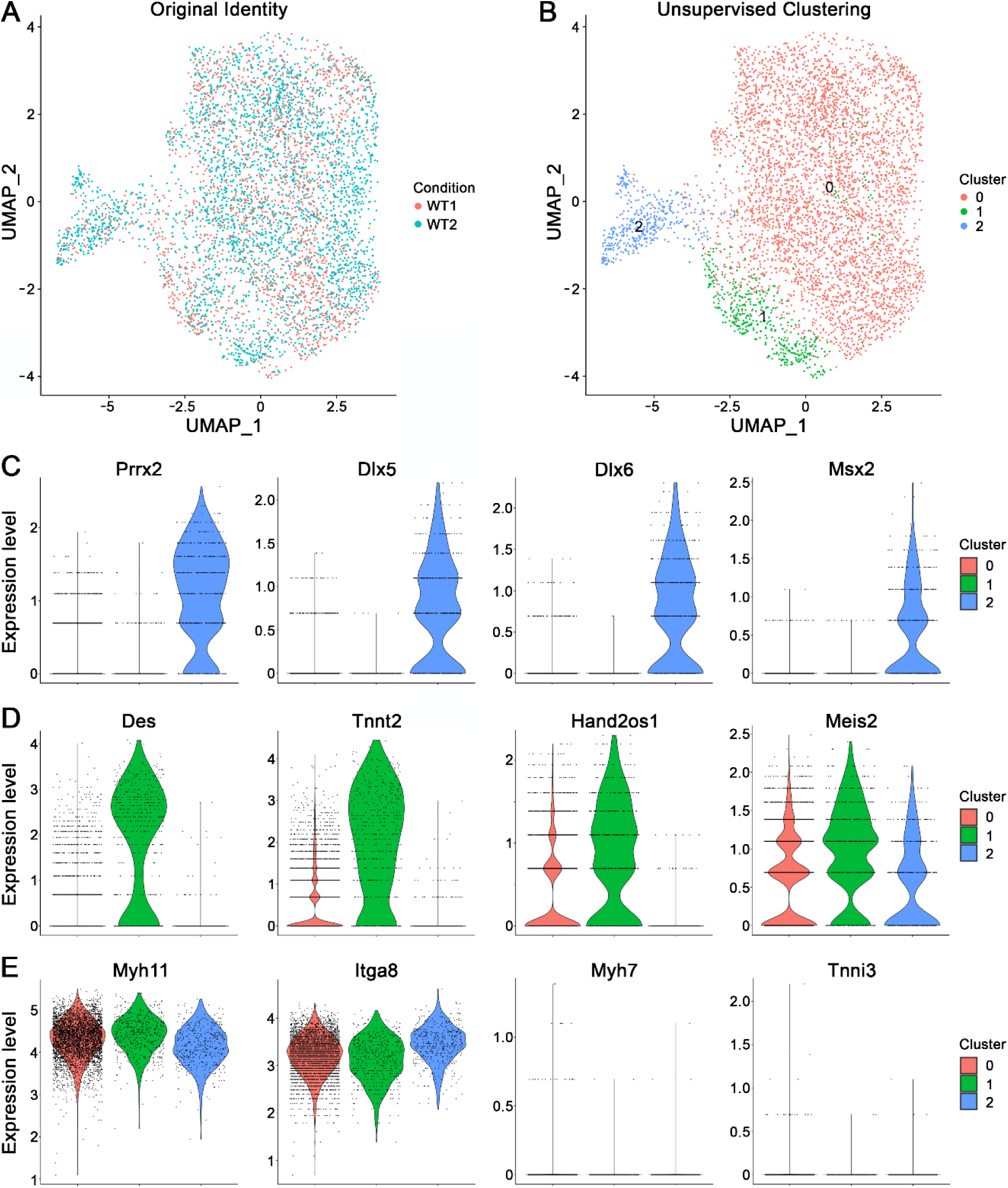
scRNA-seq of Myh11 lineage-marked SMCs of thoracic aorta. GFP+ SMCs from root, ascending, and arch aortic segments of tamoxifen-treated, 12-week-old Myh11-CreER;mT/mG (WT) mice were analyzed by scRNA-seq. (**A**) Original identity of 2 replicates and (**B**) unsupervised clustering of SMCs into 3 clusters. Expression of (**C**) neural crest-related transcription factors, *Prrx2*, *Dlx5*, *Dlx6*, and *Msx2* with maximal levels in Cluster 2, (**D**) cardiac-related markers, *Des*, *Tnnt2*, *Hand2os1*, and *Meis2* with maximal levels in Cluster 1, and (**E**) smooth muscle markers, *Myh11* and *Itga8* abundantly expressed in all 3 clusters and typical cardiomyocyte genes, *Myh7* and *Tnni3* negligibly detected in all 3 clusters.

### Reciprocal Distribution of Cardiac- and Neural Crest-Derived SMCs in the Thoracic Aorta

To identify SMCs of cardiac and neural crest origin, we used Nkx2-5-Cre and Wnt1-Cre2 mice, respectively. Crossbreeding to mT/mG strains allowed for identification of Cre-recombined GFP+ versus unrecombined RFP+ cells. Fluorescence microscopy confirmed previously described SMC distribution (4,5). Cardiac-derived SMCs constituted most of the root and the outer half of the ascending media, whereas neural crest-derived SMCs constituted all the arch and the inner half of the ascending media (Figure 2A-D). Embryological seams, however, did not correlate exactly with anatomical boundaries of aortic segments as cardiac-derived SMCs extended to the pericardial reflection rather than the origin of the brachiocephalic artery, while neural crest-derived SMCs ended before the origin of the subclavian artery on the outer curvature but after the insertion of the ligamentum arteriosum on the inner curvature rather than after the origin of the left subclavian artery (Supplemental Figure 2A-C). Other variations included scattered neural crest-derived SMCs in the aortic root and more irregular distribution of cardiac- and neural crest-derived SMCs in the lesser curvature of the ascending aorta (Supplemental Figure 2D,E). Recombination was not 100% efficient as some RFP+ medial cells were detected in root, ascending, and arch segments of compound Nkx2-5-Cre;Wnt1-Cre2;mT/mG mice, although SMCs of other embryological origins cannot be excluded (Supplemental Figure 2F,G). In summary, SMCs are mostly cardiac-derived in the aortic root, approximately equal cardiac- and neural crest-derived in the ascending aorta, only neural crest-derived in the aortic arch, and neither cardiac-nor neural crest-derived in the descending thoracic aorta.

**Figure 2:**
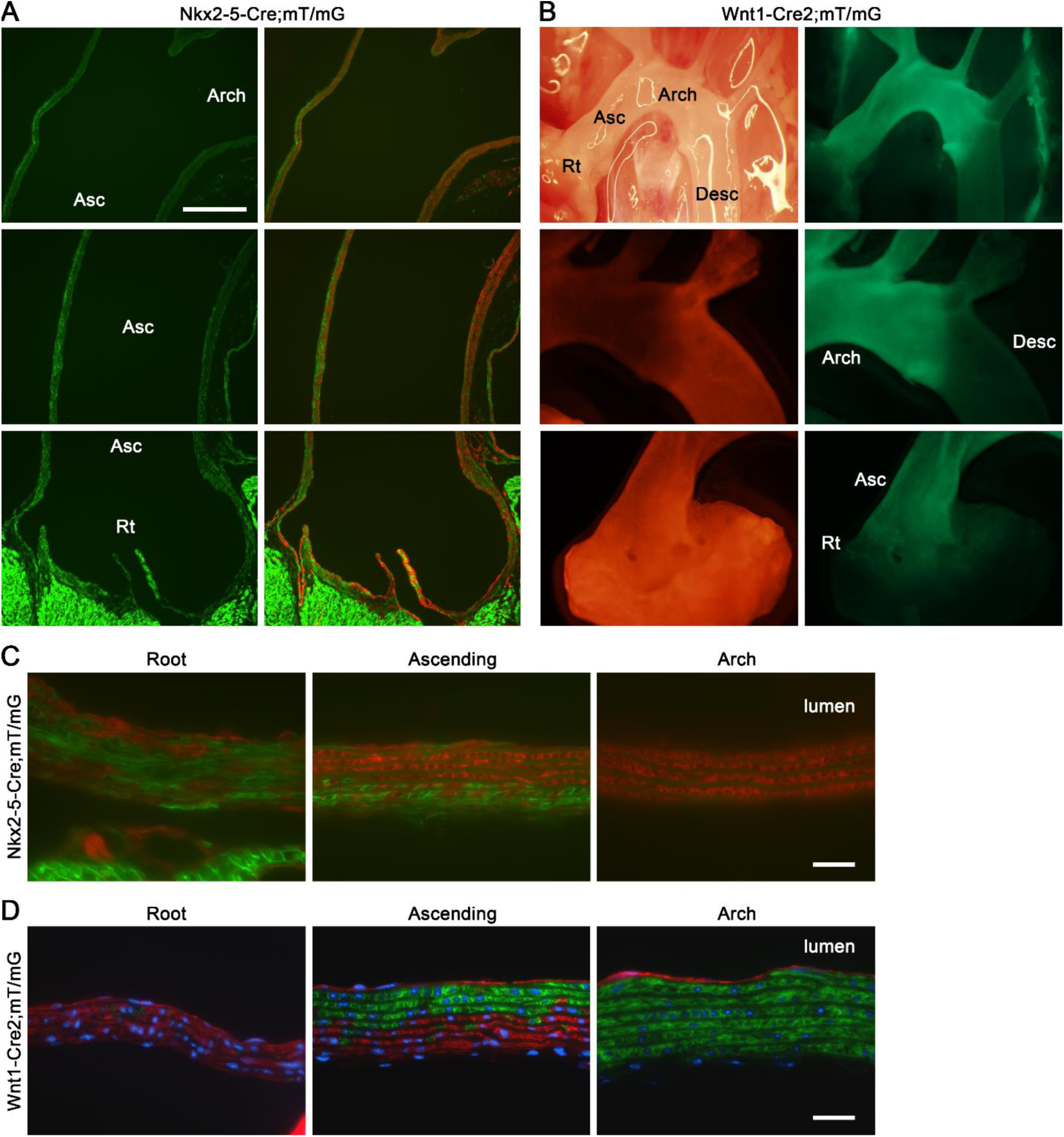
Reciprocal distribution of cardiac- and neural crest-derived SMCs in thoracic aorta. Root (Rt), ascending (Asc), arch, and descending (Desc) aortic segments were analyzed by stereo and fluorescence microscopy in reporter mice at 3 weeks of age. (**A**) GFP (bright green distinguished from dull green of elastin autofluorescence; left panels) and GFP/RFP (green and red, respectively; right panels) in longitudinal aortic sections of Nkx2-5-Cre;mT/mG mice; scale bar = 250 μm. (**B**) Light (upper left panel), RFP (lower left panels) and GFP (right panels) analysis in situ (upper row) or ex vivo (middle and lower rows) of thoracic aorta from Wnt1-Cre2;mT/mG mice. (**C**) GFP and RFP localization in longitudinal aortic sections of Nkx2-5-Cre;mT/mG mice and (**D**) GFP and RFP localization with DAPI (blue) staining of nuclei in longitudinal aortic sections of Wnt1-Cre2;mT/mG mice; orientation with intima above and adventitia below, scale bars = 25 μm.

### Heterogeneity of SMCs in Different Aortic Segments from Complementary Reporter Mice

To determine the transcriptional profile of cardiac- and neural crest-derived SMCs, we performed scRNA-seq analysis of Nkx2-5 and Wnt1 lineage-marked cells from thoracic aortas of 12-week-old mice. We separately analyzed cells from 4 segments of root, ascending, arch, and descending thoracic aorta by dividing specimens at anatomical boundaries and pooling tissue from 4-5 animals to yield sufficient cells. Limited descending thoracic aorta was processed as that segment does not contain cells of either lineage. Male Nkx2-5-Cre;mT/mG mice and both male and female Wnt1-Cre2;mT/mG mice were analyzed to assess for sex differences. In aortic segments containing SMCs of dual origin (root and ascending), the cells were separately sorted by GFP and RFP expression – except where numbers of GFP+ cells were insufficient (root segment from Wnt1-Cre2;mT/mG mice). SMCs were selected by integrin α8 expression and small numbers of dual GFP/RFP+ cells were excluded, but doublets of RFP+ SMCs with other cell types may have been included for RNA sequencing (Supplemental Figure 3A-D). Thus, a minority of cells expressing selected markers for fibroblasts (*Lum*, *Dcn*), endothelial cells (*Cdh5*, *Pecam1*), or leukocytes (*Ptprc*, *Fcer1g*) were excluded from analysis; SMC clustering was similar when other cell types were not excluded (Supplemental Figure 4A,B). Combined cell sorting and computational exclusion yielded >99.9% SMC enrichment defined as cells expressing *Myh11*. A total of 10 samples were analyzed yielding 38,717 cells meeting quality control parameters consisting of 66% Nkx2-5 vs. 34% Wnt1 lineage-marked SMCs and 20% root, 44% ascending, 34% arch, and 3% descending segment SMCs (Supplemental Figure 4C-E). The transgenes, *eGFP* and *tdTomato* were also excluded from analysis as they markedly skewed clustering. Unsupervised clustering revealed 8 SMC populations (Figure 3A). Both reporter strains contributed to each cluster and each aortic segment contributed to multiple clusters, except for a single descending aorta cluster from only Nkx2-5-Cre;mT/mG mice (descending segment cells from Wnt1-Cre2;mT/mG mice were excluded to avoid excessive SMCs not of cardiac nor neural crest origin) (Figure 3B,C). As expected, the 2 reporter strains had complementary distribution patterns with GFP+ cells from Nkx2-5-Cre;mT/mG mice overlapping with RFP+ cells from Wnt1-Cre2;mT/mG mice and vice versa (Figure 3D). Sex contributed minor differences to clustering with an X chromosome gene, *Xist* (preferentially expressed in female cells) showing less uniform distribution than that of a Y chromosome gene, *Uty* (expressed in male cells) but was not studied further in the absence of unique clusters (Supplemental Figure 4F). Together, the combined data from dual reporter mice and multiple aortic segments enable comprehensive comparison of differentially expressed genes in SMCs of different embryological origins.

**Figure 3:**
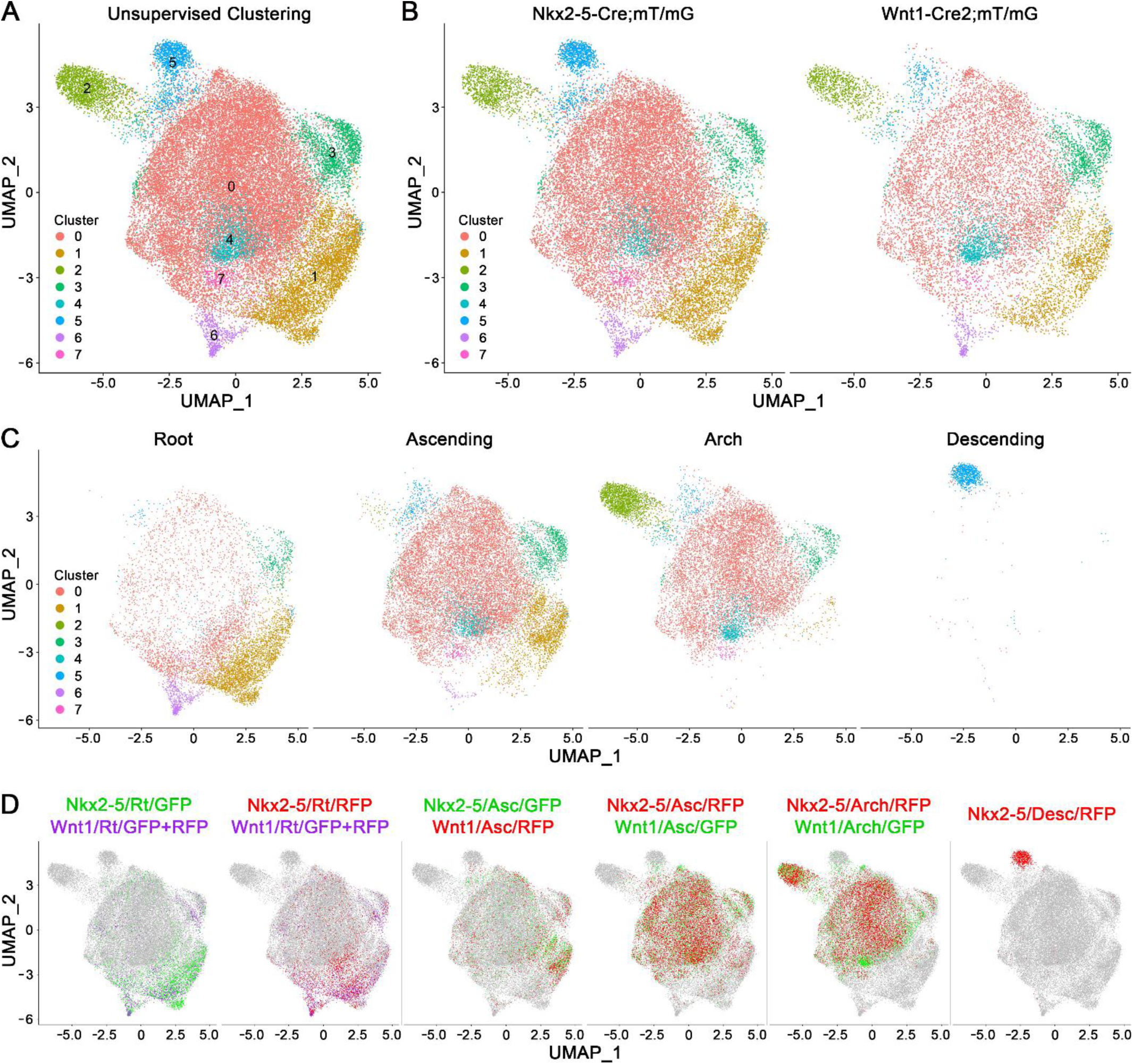
Heterogeneity of cardiac- and neural crest-derived SMCs. Integrin α8+ SMCs from root (Rt), ascending (Asc), arch, and descending (Desc) aortic segments of 12-week-old Nkx2-5-Cre;mT/mG (Nkx2-5) and Wnt1-Cre2;mT/mG (Wnt1) mice were analyzed by scRNA-seq. (**A**) Unsupervised clustering of SMCs into 8 clusters. Separation of experimental conditions contributing to clusters by: (**B**) 2 reporter strains, (**C**) 4 aortic segments, and (**D**) 10 combinations of reporter strain, aortic segment, and GFP+ or RFP+ cells.

### Cardiac-Derived SMCs Contribute to Common and Unique Clusters

Differential gene expression among cardiac-derived SMCs was examined, considering that lineage tracing by positive identification (Cre-induced GFP expression in relevant reporter) is more rigorous than negative identification (basal RFP expression in reciprocal reporter). GFP+ SMCs from Nkx2-5-Cre;mT/mG mice contributed to the largest cluster (#0) as well as a secondary cluster (#1), predominantly root cells in the latter and ascending cells in the former (Figure 4A,B). Cluster 0 consisted of both cardiac- and neural crest-derived cells from root, ascending, and arch segments and is referred to as the “common cluster” (Figure 4C). In contrast, Cluster 1 was predominantly of cardiac origin, had no arch cells, and is referred to as the “cardiac subset cluster” (Figure 4D). A heatmap of top cluster markers revealed that common Cluster 0 was characterized by genes with modest expression differences (including the top 2 markers, *Rbp4* and *Mylk4* with <2-fold change), while cardiac subset Cluster 1 had markers with greater expression differences (including the top 2 markers of *Des* and *Tnnt2* with >5-fold change) (Figure 4E). Although both were markers of cardiac-derived SMCs, *Des* predominated in ascending cells and *Tnnt2* in root cells with only partial overlap (Figure 4F,G, Supplemental Figure 5A,B). Contamination of cardiac subset Cluster 1 SMCs by cardiomyocyte RNA was unlikely as many other cardiac-specific transcripts were minimally detected (Supplemental Figure 5C). Interestingly, *Nkx2-5*, used for the reporter construct, was minimally detected, the cardiac-related transcription factors, *Gata4* and *Isl1* were detected at low abundance in both cardiac- and neural crest-derived SMCs, and *Mef2c*, an alternative second heart field reporter, was ubiquitously detected (Supplemental Figure 5D). Root cells from Wnt1-Cre2;mT/mG mice (which had not been separated by GFP and RFP fluorescence) were further examined by including reporter transgene expression and gene co-expression analysis showed strong correlation of *tdTomato*, but inverse correlation of *eGFP*, to cardiac-related genes (Supplemental Figure 5E). In summary, cardiac-derived SMCs contribute to common and unique clusters and, conversely, *Des* and/or *Tnnt2*-expressing cells constituting the cardiac subset cluster do not represent all cells of cardiac origin.

**Figure 4:**
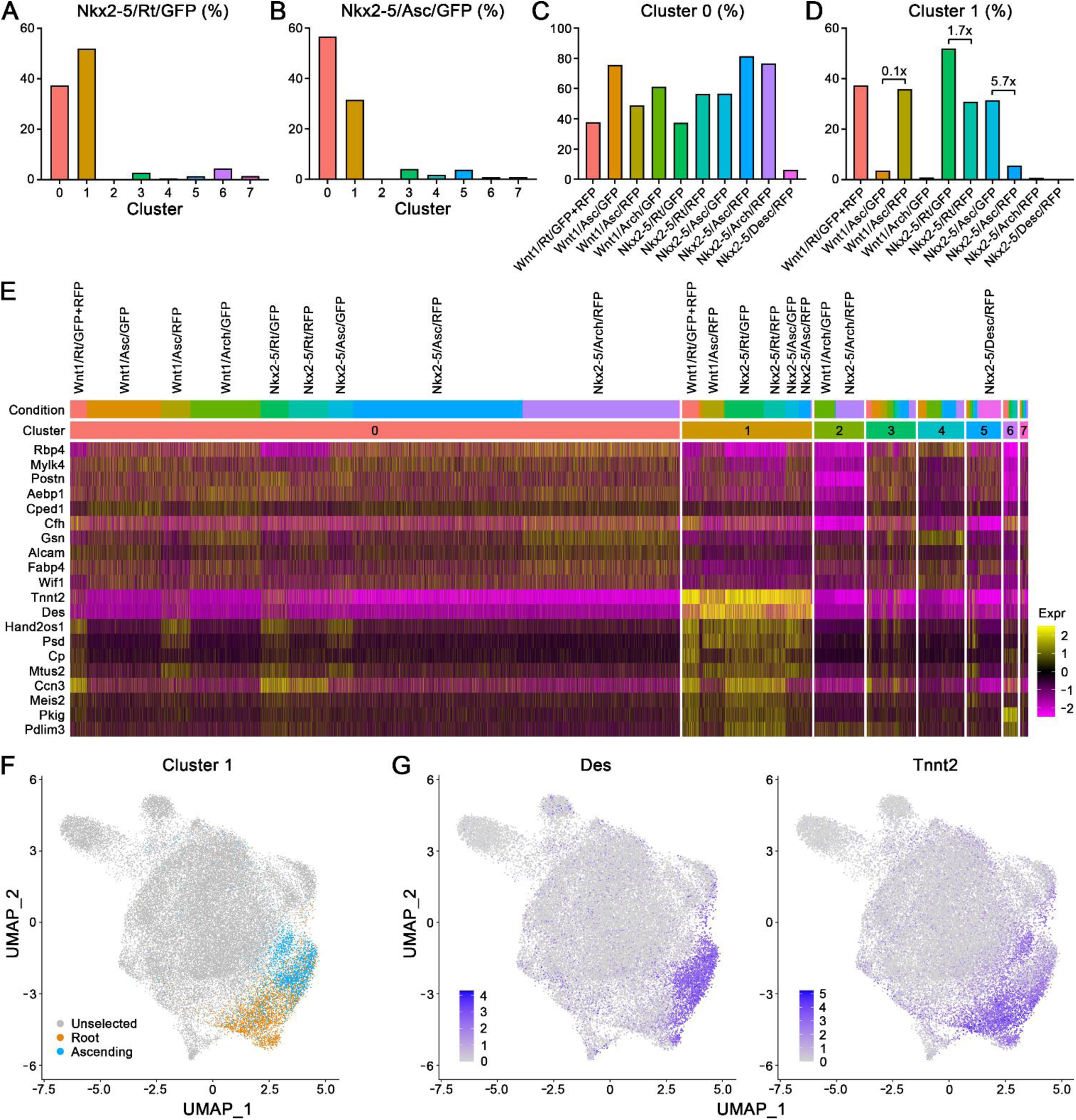
Cardiac-derived SMCs contribute to common and unique clusters. Integrin α8+ SMCs from root (Rt), ascending (Asc), arch, and descending (Desc) aortic segments of 12-week-old Nkx2-5-Cre;mT/mG (Nkx2-5) and Wnt1-Cre2;mT/mG (Wnt1) mice were analyzed by scRNA-seq to determine clustering and differential gene expression by cells of cardiac origin. Positively identified cardiac-derived cells classified as (**A**) Nkx2-5/Rt/GFP and (**B**) Nkx2-5/Asc/GFP contributed largely to Clusters 0 and 1. In turn, (**C**) Cluster 0 consisted of SMCs from all aortic segments and both reporter strains, except descending aorta, while (**D**) Cluster 1 consisted of SMCs mostly from root and ascending aorta with cells of cardiac origin predominating as indicated by fold change between paired GFP and RFP populations. (**E**) Top 10 markers for Clusters 0 and 1 by cluster and condition. (**F**) Cardiac subset cluster 1 constituents identified by origin from root and ascending segments. (**G**) Feature plots of *Des* and *Tnnt2* showing relatively restricted expression to cardiac subset Cluster 1 with preferential expression in root cells by the former and ascending cells by the latter resulting in only partial overlap.

### Neural Crest-Derived SMCs Contribute to Common and Unique Clusters

Differential gene expression among neural crest-derived SMC clusters was also examined. The majority of GFP+ SMCs from Wnt1-Cre2;mT/mG mice contributed to the largest cluster (#0), while a subset of cells from arch but not ascending segments contributed to a secondary cluster (#2) (Figure 5A,B). Whereas common Cluster 0 consisted of cells of both cardiac and neural crest origin from multiple segments, Cluster 2 consisted only of arch neural crest-derived cells and is referred to as the “neural crest subset cluster” (Figure 5C,D). A heatmap of top cluster markers revealed that neural crest subset Cluster 2 was characterized by genes with large expression differences (including *Rgs5* with >10-fold change) and/or high specificity (including *Adamtsl1* with >4-fold change as well as multiple neural crest-related transcription factors of *Prrx2*, *Dlx5*, *Dlx6*, and *Msx2* with >2-fold change) (Figure 5E). Feature plots highlighted that *Adamtsl1* and *Dlx5* were largely restricted to neural crest-derived SMCs of the aortic arch (Figure 5F,G). The neural crest-related transcription factors were not detected in other clusters, including Cluster 3 characterized by immediate early genes, Cluster 4 characterized by cytoskeletal genes, Cluster 5 characterized by microvascular and descending aorta SMC markers, Cluster 6 characterized by microvascular SMC and cardiac markers, and Cluster 7 characterized by cell cycle/cell survival genes (Supplemental Figure 6A-C). In contrast, *Rgs5* was also detected in descending aorta SMCs and certain microvascular SMCs. *Wnt1*, used for the germline reporter construct, was not detected. In summary, neural crest-derived cells contribute to both common and unique clusters and, conversely, *Adamtsl1* and *Dlx5*-expressing cells constituting the neural crest subset cluster do not represent all cells of neural crest origin.

**Figure 5:**
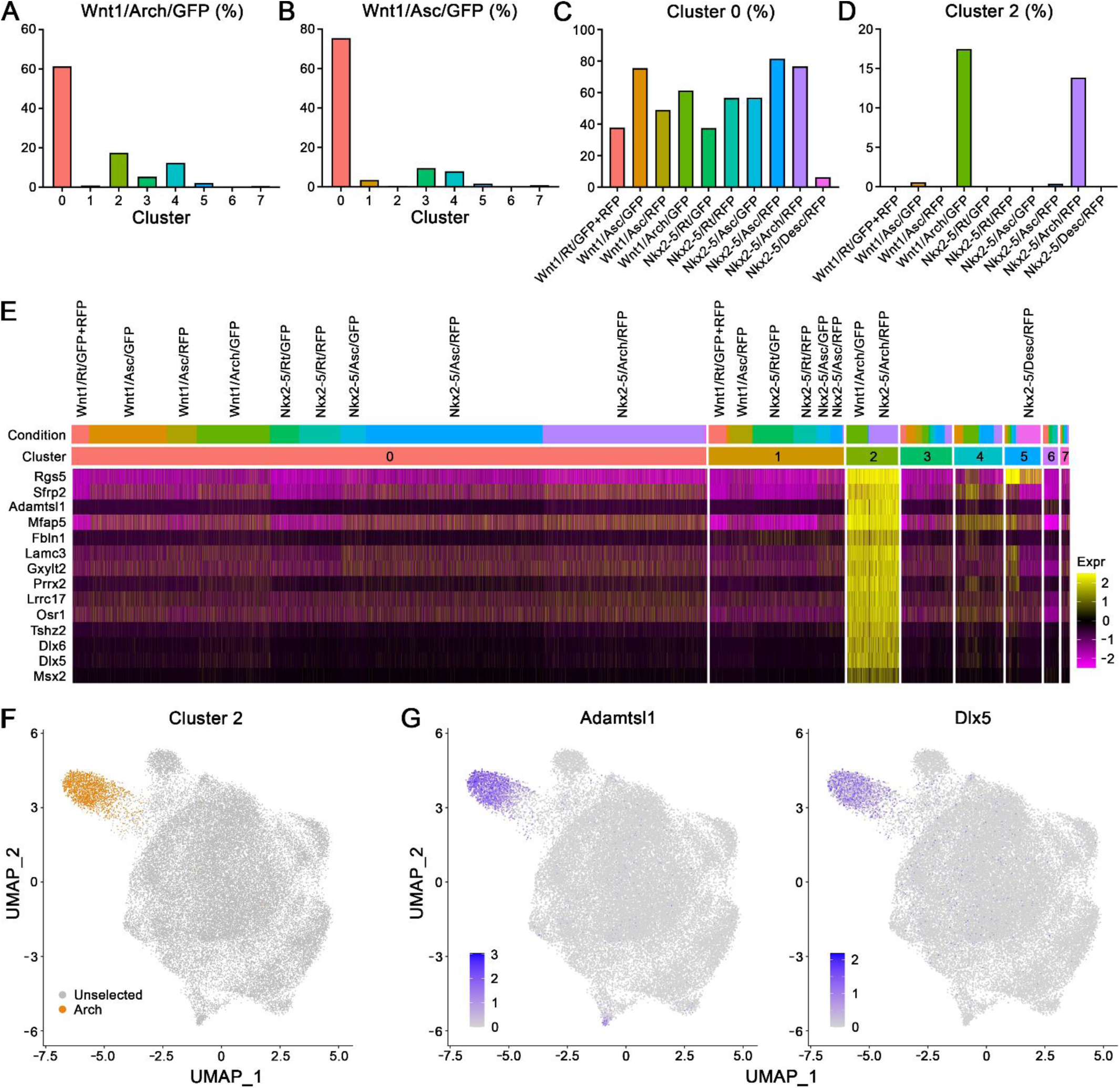
Neural crest-derived SMCs contribute to common and unique clusters. Integrin α8+ SMCs from root (Rt), ascending (Asc), arch, and descending (Desc) aortic segments of 12-week-old Nkx2-5-Cre;mT/mG (Nkx2-5) and Wnt1-Cre2;mT/mG (Wnt1) mice were analyzed by scRNA-seq to determine clustering and differential gene expression by cells of neural crest origin. Positively identified neural crest-derived cells classified as (**A**) Wnt1/Arch/GFP and (**B**) Wnt1/Asc/GFP contributed primarily to Cluster 0 with a secondary contribution to Cluster 2. (**C**) Cluster 0 consisted of both cardiac- and neural crest-derived SMCs from arch, ascending, and root segments, whereas (**D**) Cluster 2 consisted almost exclusively of neural crest-derived SMCs from the aortic arch alone. (**E**) Top 10 markers for Cluster 2, as well as 4 selected transcription factors associated with neural crest development, by cluster and condition. (**F**) Neural crest subset Cluster 2 constituents identified by origin from arch segment. (**G**) Feature plots of *Adamtsl1* and *Dlx5* showing highly restricted expression to neural crest subset Cluster 2.

### Spatial Distribution of Cardiac and Neural Crest Subset Clusters

The location of SMCs expressing markers of cardiac or neural crest subset clusters was examined. By immunofluorescence microscopy, SMCs expressing desmin were found in root and ascending segments, more commonly in the greater curvature, whereas troponin T2 was not detected in SMCs (Figure 6A). Desmin+ SMCs were also found in the pulmonary trunk and ligamentum arteriosum, including its insertion site into the proximal descending aorta. Myocardium adjacent to the aortic root provided an internal positive control for desmin and troponin T2. Reliable immunoreactivity, however, was not obtained for markers of the neural crest subset cluster. We also examined for selected transcript expression by in situ hybridization. *Des* and *Tnnt2* were detected in SMCs of root and proximal ascending segments, more in the greater curvature of the ascending aorta and the opposite curvature of the aortic root, respectively (Figure 6B). Additionally, *Dlx5* and *Msx2* were detected in the aortic arch, mostly the distal inner curvature (Figure 6C). *Dlx6* had similar distribution to *Dlx5* (not shown). These data establish that SMCs of cardiac and neural crest subset clusters localize to specific regions of thoracic aorta segments rather than uniform distribution through Nkx2-5-Cre- and Wnt1-Cre2-labeled aorta compartments.

**Figure 6:**
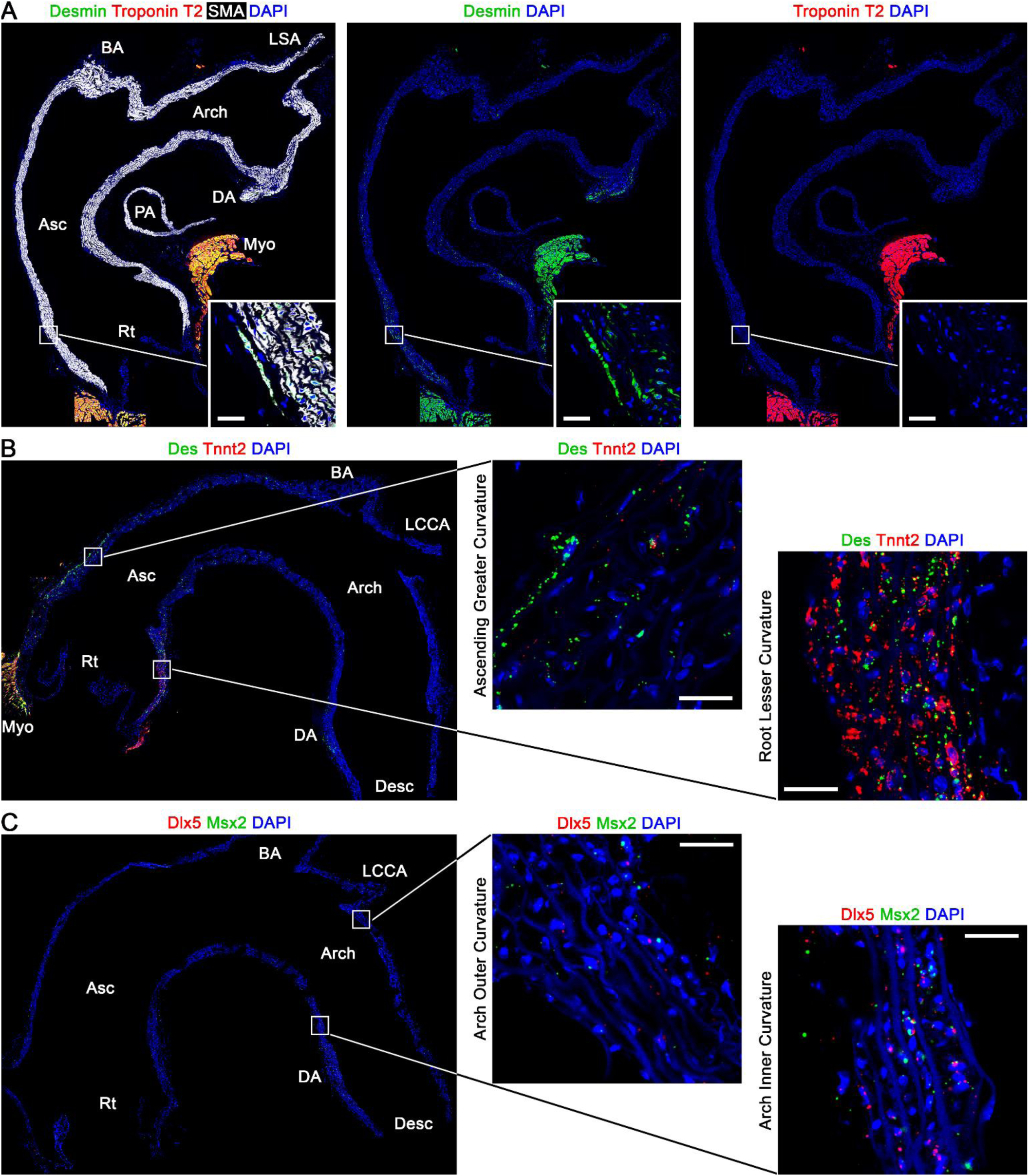
Localization of cardiac and neural crest subset SMCs. Thoracic aortas of 12-week-old C57BL/6J mice were analyzed by immunofluorescence microscopy and in situ hybridization. Detection of (**A**) desmin (green), troponin T2 (red), and smooth muscle α-actin (SMA, white) protein, (**B**) *Des* (green) and *Tnnt2* (red) transcripts, and (**C**) *Dlx5* (red) and *Msx2* (green) transcripts; nuclei labeled with DAPI (blue). Fluorescence signal in SMCs is lower for neural crest-related transcription factors than cardiac-related molecules. Selected areas of the vessel wall are magnified in insets; scale bars = 25 μm. Root: Rt, ascending: Asc, descending: desc, brachiocephalic artery: BA, left common carotid artery: LCCA, left subclavian artery: LSA, ductus arteriosum: DA, pulmonary artery: PA, and myocardium: Myo.

### Similar Transcriptional Changes after TGFβ Signaling Disruption Among Cardiac- and Neural Crest-Derived SMCs

To determine if SMCs of cardiac and neural crest origin differ in biological responses, we assessed TGFβ-dependent transcription given its critical role in aortic homeostasis and disease (10). TGFβ signaling was disrupted by deleting both type I and II receptors in 11-week-old male Tgfbr1^f/f^/Tgfbr2^f/f^;Myh11-CreER;mT/mG mice and scRNA-seq was performed of 7,373 SMCs from the aortic root, ascending aorta, and aortic arch at 12 weeks of age (Supplemental Figure 7A-C). Analysis of replicate animals was performed in the same batch with that of control Myh11-CreER;mT/mG mice described above. Unsupervised clustering revealed almost complete separation of SMC populations by TGFβ responsiveness (Figure 7A). Each population, with or without TGFβ activity, further subdivided into common, cardiac subset, and neural crest subset clusters, identified by *Rbp4* (clusters #0 and 1), *Des* (clusters #2 and 6), and *Dlx5* (clusters #3 and 5), respectively, as well as other minor clusters characterized by activation, cell cycle, and fibroblast markers (Figure 7B,C and Supplemental Figure 7D,E). Most differentially expressed genes after TGFβ signaling disruption (47 of top 50) were downregulated (Supplemental Figure 8A) and gene ontology analysis revealed marked enrichment of terms related to extracellular matrix (ECM) and TGFβ signaling regulation (Figure 7D). Notably, transcriptional changes after TGFβ signaling disruption were similar among SMCs of common, cardiac subset, and neural crest subset clusters (Supplemental Figure 8B). Although the basal expression of some TGFβ-dependent genes, e.g., *Mfap4*, *Col3a1*, *Ccn2*, and *Smad7*, was similar among SMC clusters, certain transcripts, e.g., *Eln*, *Col15a1*, *Lox*, and *Ltbp3*, were more abundant in SMCs of the neural crest subset cluster prior to receptor deletion suggesting specific TGFβ responsiveness or, alternatively, variant ECM synthesis (Figure 7E). The latter finding of skewed basal expression for certain TGFβ-dependent genes was not associated with differential expression of TGFβ ligands, receptors, and signaling effectors (Supplemental Figure 8C). Furthermore, loss of TGFβ activity did not induce changes in TGFβ ligands, receptors, and signaling effectors or angiotensin II receptors. These results indicate qualitatively similar TGFβ responses in SMCs of cardiac and neural crest origin with a quantitative difference in basal expression for a subgroup of TGFβ-dependent genes in neural crest-derived SMCs restricted to the aortic arch.

**Figure 7:**
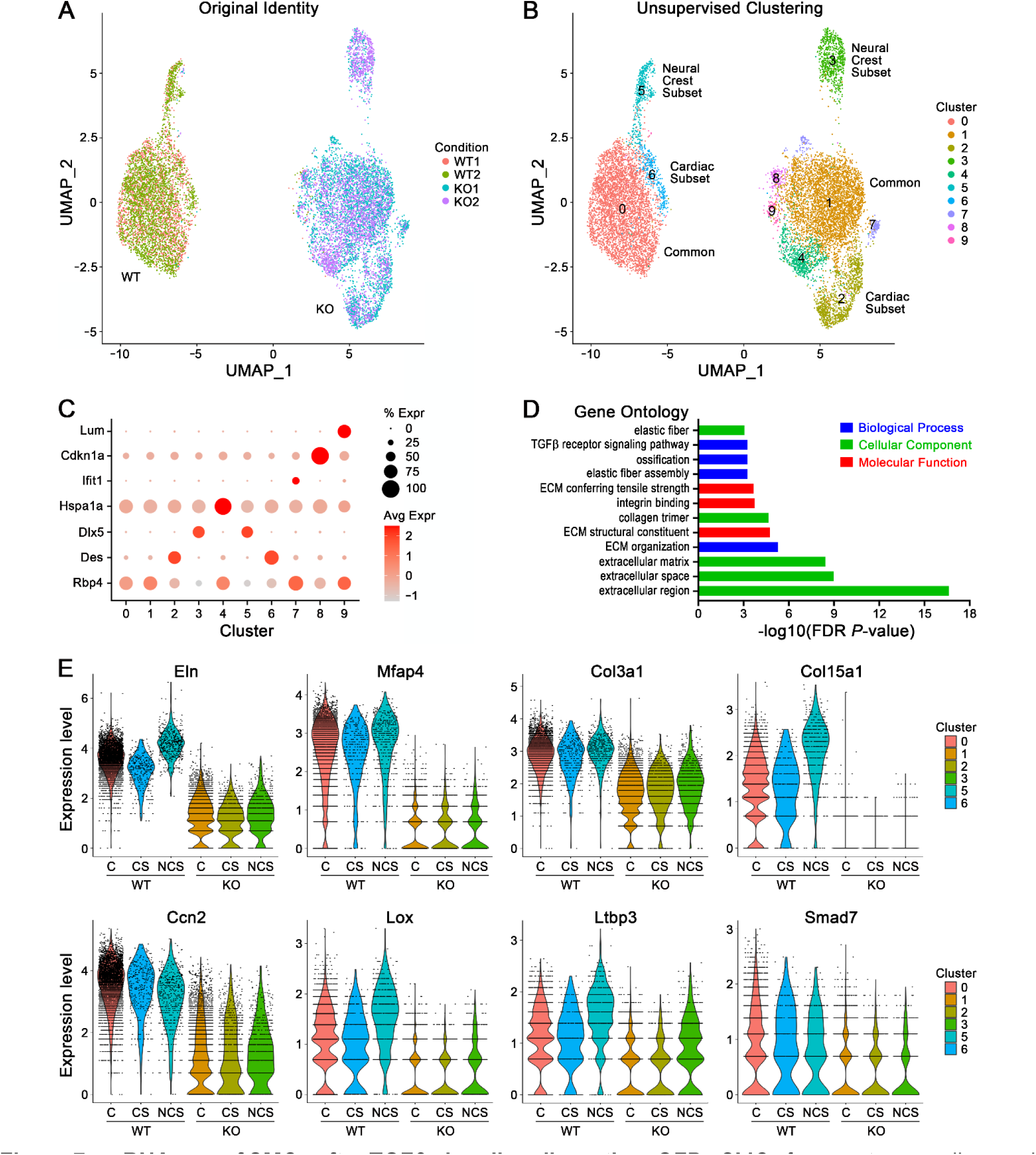
scRNA-seq of SMCs after TGFβ signaling disruption. GFP+ SMCs from root, ascending, and arch aortic segments of tamoxifen-treated, 12-week-old Myh11-CreER;mT/mG (WT) and Tgfbr1^f/f^/Tgfbr2^f/f^;Myh11-CreER;mT/mG (KO) mice were analyzed by scRNA-seq. (**A**) Original identity by strain and (**B**) unsupervised clustering into 10 clusters. (**C**) Percent and average (avg) expression (expr) of cluster markers, including *Rbp4* to identify common (#0 and 1), *Des* to identify cardiac subset (#2 and 6), and *Dlx5* to identify neural crest subset (#3 and 5) clusters. (**D**) Gene ontology enrichment analysis of top 50 differentially expressed genes for KO vs. WT cells. (**E**) Expression of selected ECM (*Eln*, *Mfap4*, *Col3a1*, *Col15a1*, *Ccn2*, and *Lox*) and TGFβ signaling modulator (*Ltbp3* and *Smad7*) genes in common (C), cardiac subset (CS), and neural crest subset (NCS) clusters of WT and KO cells.

### Few Markers of Cardiac and Neural Crest Subset Clusters are Detected in Human Aortas

We investigated if our findings in mice are relevant for humans. Root, ascending, arch, and descending aorta specimens were obtained from adult organ donors without aortic disease, though isolation of vessel wall cells was problematic because of tenacious ECM and a lack of reagents to select SMCs. Despite greater than 1,000-fold tissue mass, rapid processing of specimens, removal of adventitia, enzymatic digestion at 37 °C, exclusion of leukocytes and erythrocytes, and greater sequencing depth, the yield of 3,851 vessel wall cells was less than that of murine aortas with fewer genes and total transcripts detected per cell (Supplemental Figure 9A-C). Unsupervised clustering revealed several clusters of SMCs/pericytes, fibroblasts, and endothelial cells (Figure 8A,B). SMC/pericyte clusters #2, 4, and 5 did not segregate by aortic segment but were characterized by microvascular, macrovascular, and activation markers, respectively (Supplemental Figure 9D). Markers of murine cardiac- and neural crest-derived SMCs, many detected at low levels, were not differentially expressed by human SMCs/pericytes, except for increased *RGS5* and *DLX5* in microvascular SMCs/pericytes of Cluster 2 and activated SMCs/pericytes of Cluster 5 (Figure 8C). *RGS5* and *DLX5*, however, were not differentially expressed by macrovascular SMCs of Cluster 4 from different aortic segments to correlate with diverse embryological origins (Figure 8D). In a second subject, fewer SMCs/pericytes were obtained yielding a single cluster with fibroblasts as leukocytes were markedly over-represented after analysis of all isolated cells from specimens that included the adventitia earlier in the development of our technique (Supplemental Figure 9E,F). To avoid problems from cell isolation, we performed bulk RNA-seq of aortic tissue from 4 segments of 5 individuals but 2 arch and 3 descending specimens were not sequenced because of insufficient RNA quality. Aortic root non-coronary sinus tissue was characterized by several cardiac-related genes, e.g., *TCF21*, *NKX2-5*, *DES*, *ACTA1*, *MEIS2*, and *GATA4*, mid-ascending tissue at the greater curvature by proximal HOX genes, e.g., *HOXA3*, *HOXA4*, *HOXA2*, and *HOXA1*, inner mid-arch tissue by diverse cell fate determining genes, e.g., *MAB21L2*, *MAB21L1*, *SOX6*, *DLX1*, and *TFAP2A*, and proximal descending thoracic tissue by more distal HOX genes, e.g., *HOXB7*, *HOXB6*, and *HOXD8* (Figure 8E). Of many cardiac and neural crest markers identified in mouse aortas, only *DES* and *MEIS2* were differentially expressed in human aortic segments though restricted to the root and there were few differences between mid-ascending and inner arch tissue suggesting a more proximal transition between SMC lineages. Although species, age, and/or specimen location differences may account for the discrepant findings to murine SMCs, we cannot exclude technical limitations in analyses of human aortas.

**Figure 8:**
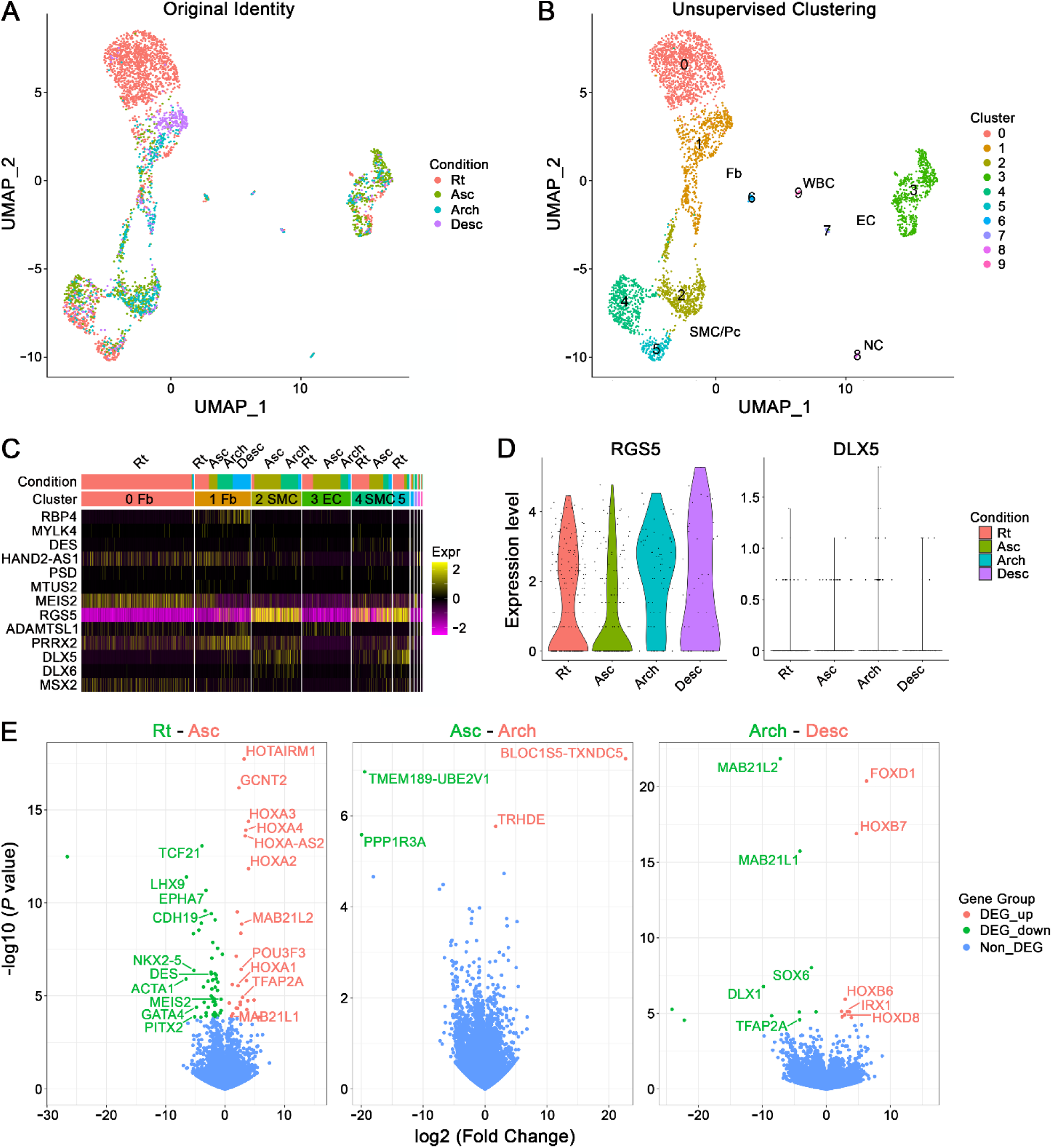
scRNA-seq of human aortic SMCs. CD45-/CD235a-vessel wall cells from root (Rt), ascending (Asc), arch, and descending (Desc) aortic segments of 55-year-old, male organ donor were analyzed by scRNA-seq. (**A**) Original identity by aortic segment and (**B**) unsupervised clustering into 10 clusters of SMCs/pericytes (Pc; #2, 4, and 5), fibroblasts (Fb; #0, 1, and 6), endothelial cells (EC; #3 and 7), neuronal cells (NC; #8), and leukocytes (WBC; #9). (**C**) Relative expression of murine common, cardiac subset, and neural crest subset markers by cluster and aortic segment (*TNNT2* was not detected in human vessel wall cells). (**D**) Expression levels of *RGS5* and *DLX5* in macrovascular SMCs of Cluster 4 by aortic segment. Additionally, aortic specimens with adequate RNA quality from root (*n* = 5), ascending (*n* = 5), arch (*n* = 3), and descending (*n* = 2) segments of 5 organ donors were analyzed by bulk RNA-seq. (**E**) Volcano plots of differentially expressed genes (DEG) with adjusted *P* values <0.05 and fold change <-2 (green) or >2 (orange) for comparisons between ascending vs. root (*n* = 80 DEG), arch vs. ascending (*n* = 4 DEG), and descending vs. arch (*n* = 19 DEG) aortic segments.

## Discussion

scRNA-seq has been widely applied to characterize vessel wall cells among different organs in mice (21). Despite providing unprecedented insight into the heterogeneity of diverse vascular beds, broadly designed studies are limited in defining more subtle differences within a blood vessel. Analysis of several hundred aortic SMCs, among thousands of cells, identified differential gene expression along the aorta, e.g., *Des* in the proximal aorta and *Prrx2* in the distal aorta, yet distinct cell clusters by embryological origin or aortic segment were not defined (22). We interrogate a single cell type (SMC) from a single blood vessel (thoracic aorta) to power comparisons of gene expression differences. Previous scRNA-seq analyses of mature thoracic aorta cells have identified differential expression of cardiac- and neural crest-related genes, such as *Des*, *Tnnt2*, *Rgs5*, *Prrx2*, *Dlx5*, *Dlx6*, and *Msx2* (23–25). Without dual lineage tracing, however, clusters with diverse embryological origins were not apparent and, without separation of aortic segments, localization of clusters was not evident. We find multiple subtypes of cardiac- and neural crest-derived SMCs in the thoracic aorta of adult mice. Most cells constitute a common cluster not distinguishable by embryological origin, while a minority contribute to distinct cardiac and neural crest subset clusters. Our dual reporter, multi-segment data provide a resource to identify embryological origins of mature aortic SMCs, to define lineage subtypes and regulatory states, and to inform conditional Cre deletion systems applicable to adult animals.

The common cluster is characterized by markers with relatively low expression differences and specificity, such as *Rbp4* and *Mylk4* previously reported as markers of aortic SMCs versus other aortic and cardiac mural cells (22). Different transcriptional profiles of common cluster SMCs versus cardiac and neural crest subset SMCs imply that progenitors differentiate to various lineage subtypes or immature cell types undergo different postnatal transitions. Indeed, several types of outflow tract and neural crest-derived SMCs with distinct molecular signatures are found during embryogenesis, and genomic profiles markedly change during postnatal maturation of aortas (26–28). Previously documented differences in neural crest-derived SMCs from pharyngeal arch arteries versus outflow tract does not explain our observation that most arch, ascending, and root SMCs contribute to a common cluster (29). Further work at multiple developmental and maturation stages is necessary to determine the chronology and regulatory networks governing divergence of common from subset clusters as well as convergence of cardiac- and neural crest-derived intermediates to a common cluster.

The cardiac subset cluster markers, *Des* and *Tnnt2* have relatively high expression differences and specificity. Typical cardiac transcription factors are near undetectable or not specific for cardiac-derived SMCs, the recently described cardiac regulator, *Hand2os1* an exception (20). *Des* and its product, desmin predominate in proximal ascending SMCs corresponding to previous immunofluorescence analysis (22), while *Tnnt2* predominates in aortic root SMCs as shown by prior in situ hybridization (25). Although troponin was reported in SMCs along the ascending aorta extending into the distal aortic arch (30), our data does not endorse cardiac subset cluster cells in distal locations except at the insertion site of the ductus arteriosum. Furthermore, we were unable to detect protein immunoreactivity within any SMCs, possibly because the troponin complex is not assembled in the absence of ATPase-inhibitory and calcium-binding subunits (transcripts encoding troponin I and troponin C isoforms were not detected in SMCs). *Tnnt2* has been tracked in the developing outflow tract via reporter activity and SMC expression may represent persistence of a few cardiac-related genes following transdifferentiation from myocardial precursors (26). Mechanisms for differential expression of *Des* and *Tnnt2* among SMCs of root versus ascending segments or greater versus lesser curvatures are not known but may relate to more than one cardiac subset cluster as discussed below. Although there was no impact on SMC clustering, our strategy to diminish artifact by excluding contaminating fibroblasts, endothelial cells, and leukocytes may have excluded atypical root SMCs expressing *Dcn* and *Lum* (22,25) and we are examining if fibroblast-like SMCs are an uncommon cell type or artifact.

The neural crest subset cluster is characterized by numerous markers of varying specificity. *Rgs5* with the greatest fold-change expression difference is also expressed by microvascular and descending thoracic aortic SMCs. Other distinguishing genes, such as *Adamtsl1*, *Fbln1*, and *Prrx2*, with high specificity in our analysis of aortic SMCs alone are also expressed by arterial SMCs, fibroblasts, and endothelial cells (22). The neural crest-related homeobox transcription factors, *Dlx5*, *Dlx6*, and *Msx2* that show significant segmental differences in our data are relatively specific for distal aortic SMCs in a broader analysis of several vascular beds (22). During embryonic development, *Dlx5*, *Dlx6*, and *Msx2* are characteristic of a subtype of neural crest-derived SMCs found in cardiac arteries but not outflow tract (27), and of distal pharyngeal arch arteries (29). Additional marker genes of the neural crest subset cluster that we did not discuss because of lower (though still significant) expression changes include other transcription factors and signaling molecules related to neural crest development, such as *Sox9*, *Sema3c*, and *Ednra*. Several homeobox transcription factors, including *Dlx5*, are known to bind Smad signal transducers (31), and persistent expression may alter basal levels for a subgroup of TGFβ-dependent genes. Alternatively, endothelin enhances certain TGFβ signaling (32), and type A receptor expression may relate to skewed ECM synthesis by neural crest subset SMCs.

Categorical disruption of TGFβ signaling by conditional deletion of both type I and II receptors unequivocally identifies TGFβ-dependent genes in the absence of alternate receptor complex activity or paradoxical receptor overactivity. Gene expression changes are uniform among aortic SMCs regardless of embryological origins. Most genes are downregulated, including many ECM molecules and TGFβ signaling modulators, while a few genes are upregulated, such as *Itih4*. The loss of many ECM molecules was not evident in whole genome expression analysis of aortic tissue after TGFβ signaling disruption in SMCs, likely masked by adventitial fibroblast transcripts (10). We did not identify induction of transcripts encoding TGFβ ligands or angiotensin II receptor as previously reported in aortic SMCs after partial loss of TGFβ signaling (15). In the latter murine model with heterozygous inactivating *Tgfbr1* germline mutation, neural crest-derived SMCs remain signaling sufficient while cardiac-derived SMCs become signaling deficient.

Differences in TGFβ signaling among SMCs of different embryological origin after partial gene inactivation may reflect the greater transcription of a subset of TGFβ-dependent genes in SMCs of the neural crest subset cluster. Of note, SMCs of the neural crest subset cluster are restricted to the aortic arch and these cells are unlikely to participate in dilatation of the aortic root typical of Loeys-Dietz syndrome.

The methodology has numerous limitations. Lineage tracing is not precise depending on reporter gene specificity, construct regulatory elements, and Cre recombinase efficiency. We find <0.2% ectopic expression in descending SMCs and 1-10% incomplete expression in compound reporters. Nkx2-5-Cre signal is relatively weak in SMCs (compared to cardiomyocytes) and is not specific for first versus second heart field as all cardiac chambers are marked (34), while Wnt1-Cre2 marks both cardiac neural crest- and cranial neural crest-derived cells (35). Hence our preference for broad descriptors of cardiac-derived and neural crest-derived SMCs rather than second heart field-derived and cardiac neural crest-derived SMCs. Dividing aortic segments at anatomical boundaries does not precisely correspond to embryological seams, thus some neural crest subset cluster SMCs may be included in extra-pericardial ascending aorta or descending thoracic SMCs from the outer curvature of the distal aortic arch. The expression of many developmental regulatory genes is extinguished with maturation and identification of embryological origin in adults relies on few, generally low-abundance genes. Furthermore, the process of isolating single cells triggers artifact responses within minutes unless transcriptional arrest is induced via hypothermia (16). Given our interrogation of a single cell type, we selected a relatively low-resolution parameter (0.2) for clustering granularity to avoid conceptual complexity yet still yielding 8 SMC clusters from 3 embryological origins (including the descending thoracic segment), several regulatory states, and inadvertent inclusion of 2 microvascular SMC populations (perivascular and cardiac) from contiguous tissue – although selection by Myh11 reporter or integrin α8 expression prevented contamination with pericytes. Sequentially increasing resolution (to 1.2) does not affect cell distribution but markedly increases cluster numbers (to 22).

The common cluster segregates into several clusters, the cardiac subset cluster sequentially separates as: *Des*-expressing root and ascending SMCs, *Des*- and *Tnnt2*-expressing root SMCs, *Tnnt2*-expressing ascending SMCs, and *Tnnt2*-expressing root SMCs, while the neural crest subset cluster does not subdivide.

scRNA-seq analysis of human aortas is more challenging than in mice. The media is thicker and the ECM is stronger that limits isolation of SMCs with over-representation of other cell types, particularly leukocytes. Even the relatively few SMCs isolated are skewed by over-representation of microvascular SMCs from vasa vasorum with less tenacious ECM. Lower quality RNA and fewer genes per cell may represent the age and comorbidities of human subjects. Additionally, analysis of selected sections of aortic segments, e.g., mid-ascending or mid-arch, may exclude regions with more numerous SMCs of cardiac and neural crest subset clusters. Other scRNA-seq studies of human aortas, however, have not identified cardiac- or neural crest-derived SMC clusters (30,36,37). While certain technical limitations are avoided by bulk RNA-seq analysis of human aortas yielding more segmental markers, particularly HOX genes, the absence of lineage tracing precludes definitive association with SMC embryological origin.

In conclusion, multiple subtypes of cardiac- and neural crest-derived SMCs in adult thoracic aortas arise during embryological and/or postnatal development. We do not identify genes suitable for conditional deletion strategies targeting whole embryological lineages in adult SMCs, although *Des* and *Tnnt2* are appropriate for a subset(s) of cardiac-derived SMCs and *Dlx5*, *Dlx6*, or *Msx2* are appropriate for a subset of neural crest-derived SMCs. Aortic SMCs are uniformly impacted by disruption of TGFβ signaling, even though a subset of arch-restricted, neural crest-derived SMCs have skewed basal expression of certain TGFβ-dependent genes. While different transcriptional states do not necessarily signify different cellular phenotypes, they potentially underlie mechanisms for varying disease susceptibility along the aorta. It is not known if SMC minor clusters with vestigial gene expression or the major cluster without distinguishing embryological markers differ in pathogenicity, perhaps dependent on disease process.

## **Methods** (Expanded Methods are included in the Supplemental Materials)

### Mice

C57BL/6J (strain #000664), Nkx2-5-Cre (strain #024637), Wnt1-Cre2 (strain #022501), and mT/mG (strain #007676) mice were purchased from The Jackson Laboratory. Myh11-CreER^T2^ mice (38) were obtained from Dr. Stefan Offermanns, University of Heidelberg, Tgfbr1^f/f^ mice (39) were obtained from Dr. Martin M. Matzuk, Baylor College of Medicine, and Tgfbr2^f/f^ mice (40) were obtained from Dr. Harold L. Moses, Vanderbilt University. Mutant strains were bred to a C57BL/6J background for >6 generations. Male (C57BL/6J, Myh11-CreER;mT/mG, Nkx2-5-Cre;mT/mG, Wnt1-Cre2;mT/mG, and Tgfbr1^f/f^/Tgfbr2^f/f^;Myh11-CreER;mTmG) and female (C57BL/6J and Wnt1-Cre2;mT/mG) mice were euthanized at 3 and 12 weeks of age for thoracic aorta analyses (the bacterial artificial chromosome containing Myh11-CreER inserted into the Y chromosome in this strain and female mice do not express the construct).

### Human Subjects

Non-dilated, non-atherosclerotic aortas were procured from organ donors whose hearts were not used for clinical transplantation. scRNA-seq analyses were performed in aortic tissue from 2 subjects (51-year-old, White, non-Hispanic female and 55-year-old, White, non-Hispanic male). Bulk RNA-seq was performed in aortic tissue from 5 subjects (19-year-old, Black, non-Hispanic male, 39-year-old, Black, non-Hispanic male, 40-year-old, White, Hispanic male, 48-year-old, White, non-Hispanic male, and 60-year-old, White, non-Hispanic female).

### Study Approval

Animal research protocols were approved by the Institutional Animal Care and Use Committee of Yale University. Human subjects research protocols were approved by New England Donor Services (with signed consent for research by next-of-kin of deceased subjects) and the Institutional Review Board of Yale University (with waiver for consent).

### Statistics

Gene co-expression analysis by Pearson correlation was performed using Prism 9.2.0 (GraphPad Software). scRNA-seq analyses were performed using R (r-project.org), markers were ranked by adjusted *P* value or by fold change if *P*-values could not be differentiated. Bulk RNA-seq analyses were performed using DESeq2; adjusted *P* value <0.05 was considered significant and fold change >2 or <-2 was used as cutoff for differential expression.

## Abbreviations

ECM: extracellular matrix
GFP: green fluorescent protein
RFP: red fluorescent protein
RNA-seq: RNA sequencing
scRNA-seq: single-cell
RNA: sequencing
SMC: smooth muscle cell
TGFβ: transforming growth factor-β

