## Supplemental_Materials for "Heterogeneous Cardiac- and Neural Crest-Derived Aortic Smooth Muscle Cells have Similar Transcriptional Changes after TGFβ Signaling Disruption"

**Supplemental Materials**: expanded methods, 9 figures

**Expanded Methods**

**Animal Treatment.** Cre-Lox recombination was induced by tamoxifen (T5648, Sigma-Aldrich) at 2 mg/day i.p. for 5 days starting at 11 weeks of age in male Myh11-CreER;mTmG and Tgfbr1$^{f/f}$/Tgfbr2$^{f/f}$;Myh11-CreER;mTmG mice.

**Confocal Microscopy.** Formalin-fixed, paraffin-embedded or frozen, optimum cutting temperature (OCT) compound-embedded aortic sections were incubated with FITC anti-smooth muscle α-actin (F3777, Sigma-Aldrich), anti-desmin (5332, Cell Signaling Technology), or Alexa Fluor 647 anti-cardiac troponin T (565744, BD Biosciences) overnight at 4 °C. Unconjugated primary antibody was detected with Alexa Fluor 568 anti-rabbit IgG (A10042, Invitrogen). The sections were mounted with ProLong Gold Antifade reagent with DAPI (P36931, Life Technologies) and images were acquired with a confocal microscope (SP8, Leica).

**Cell Isolation.** Mouse aortas were minced, incubated in 0.5 ml DMEM with 1.5 mg/mL collagenase A (10103578001, Roche) , 0.5 mg/mL elastase (LS002294, Worthington Biochemical), 10 mg/mL *Bacillus licheniformis* protease (P5380, Sigma-Aldrich), 10 mg/mL Dispase II (D4693, Sigma-Aldrich), 125 U/mL DNase I (DN25, Sigma-Aldrich) and 0.2% BSA (A9418, Sigma-Aldrich) for 3 h at 4 °C and passed through a 40 μm filter (1,2). The cells were incubated with cell-impermeant viability dye (65-0865-14, Invitrogen), blocked with anti-mouse FcγRI/II/III/IV (139302,101302, and 149502, BioLegend) for 20 min at 4 °C, and stained with custom APC anti-mouse integrin α8 (same clone as AF4076, R&D Systems) for 30 min at 4 °C. The cells were pelleted and resuspended in 0.4% BSA/PBS for fluorescence-activated cell sorting. Alternatively, human aortas were minced and digested in 10 ml DMEM with 10% FBS, 1.5 mg/mL collagenase A, and 0.5 mg/ml elastase for 1 h at 37 °C and passed through a 100 μm filter. The cells were incubated with anti-human FcγRI/II/III (422302, BioLegend) for 5 min and stained with viability dye (65-0865-14, Invitrogen) for 20 min at 4 °C. To enrich for vascular cells by excluding leukocytes and erythrocytes, isolated aortic cells were labeled with PE-anti-human CD45 (304008, BioLegend) and FITC anti-human CD235a (349104, BioLegend) for 30 min at 4 °C, washed, and resuspended in 0.4% BSA/PBS for fluorescence-activated cell sorting.

**scRNA-seq.** Viable integrin α8+ SMCs (for mice) or CD45-/CD235a- vascular cells (for humans) were sorted with a FACSAria (BD Biosciences) and collected in 0.04% BSA/PBS. The selected cells were processed for scRNA-seq library preparation using the Chromium™ Single Cell Platform (10x Genomics) as per the manufacturer's protocol. Briefly, single cells were partitioned into Gel Beads in Emulsion using the Chromium™ system (10x Genomics), followed by cell lysis and barcoded reverse transcription of RNA, cDNA amplification and shearing, and 5' adaptor and sample index attachment. The single-cell RNA-seq libraries were sequenced on a Novaseq 6000 System (Illumina) at the Yale Center for Genome Analysis. For transcript analyses, the fastq data from the barcoded library sequence was processed using Cell Ranger (10x Genomics), mapped to a mouse reference genome (mm10) customized by the addition of sequences and annotations for exogenous *eGFP* and *tdTomato* genes or a human reference genome (GRCh38), and further processed in R using Seurat V4.3 (3). Data for mouse SMCS were filtered as follows: i) cells with <2,000 genes or >6,000 genes were excluded, ii) cells with <5,000 or >30,000 UMIs were excluded, iii) cells with >10% mitochondrial gene expression were excluded, and iv) transcripts *Gm42418* and *AY036118* were removed from the count matrix since they overlap an unannotated *Rn45s* rRNA locus and may include counts from rRNA molecules amplified during library preparation (4). Additional filtering in analyses from Nkx2-5-Cre and Wnt1-Cre2 mice included: v) transcripts *eGFP* and *tdTomato* were removed as they skewed clustering of reciprocally labelled lineages and vi) cells with ≥1 copy of *Dcn, Lum, Cdh5, Pecam1, Ptprc,* or *Fcer1g* were excluded to minimize doublets of SMCs with other vessel wall cell types. Data for human aortic cells (including adventitial cells) were filtered as follows: i) cells <500 or >4,000 genes were excluded, ii) cells <1,000 or >20,000 UMIs were excluded, and iii) cells with >15% mitochondrial gene expression were excluded. Alternatively, data for human aortic medial cells (excluding adventitial cells) were filtered as follows: i) cells with <1,000 or >5,000 genes were excluded, ii) cells with <1,500 or >25,000 UMIs were excluded, and iii) cells with >15% mitochondrial gene expression were excluded. The data was log normalized by SCTransform with default settings (5). For combined analyses from Nkx2-5-Cre and Wnt1-Cre2 mice, integration was used according to Seurat pipelines (6). Genes with the most variable expression were used for clustering by Uniform Manifold Approximation and Projection (UMAP) and these projections were used for visualization. The Seurat FindAllMarkers function was used to identify markers across all samples and the Seurat FindMarkers function was used to analyse markers between two groups to generate differential expression. The data has been deposited in NCBI's Gene Expression Omnibus (GEO): GSE253935 and GSE194083.

**Bulk RNA-seq.** Crushed human aortic tissue was immersed in RLT lysis buffer (79216, Qiagen) and vigorously vortexed. Total RNA was isolated using a RNeasy Mini Kit (74104, Qiagen) and DNase Digestion Set (79254, Qiagen) according to the manufacturer's protocol. Quality control was assessed by nanodrop and an Agilent Bioanalyzer. Next-generation, whole-transcriptome sequencing was performed using a NovaSeq 6000 System (Illumina) at the Yale Center for Genome Analysis. Low quality reads were trimmed, and adaptor contamination were removed using Trim Galore (v0.5.0). Trimmed reads were mapped to the human reference genome (GRCh38) using HISAT2 (v2.1.0) (7). Gene expression levels were quantified using StringTie (v1.3.3b) (8) with gene models (v27) from the GENCODE project. Differentially expressed genes were identified using DESeq2 (v 1.22.1) (9). The data has been deposited in NCBI's GEO: GSE253935.

**Gene Ontology Enrichment Analysis.** The top 50 differentially expressed genes between experimental groups by false discovery rate-adjusted *P* values (and by fold change if *P* values could not be differentiated) were used for gene ontology enrichment analysis of RNA-seq datasets. The gene list was analyzed using DAVID v2024q1 (https://david.ncifcrf.gov) to identify enriched biological themes among biological process, cellular component, and molecular function terms. Enriched terms were ranked by false discovery rate-adjusted *P*-value.

**RNA In Situ Hybridization.** Thoracic aortas from 12-week-old male C57BL/6J mice were fixed in 4% paraformaldehyde overnight at 4 °C, incubated in 15% sucrose overnight at 4 °C, and embedded in OCT (4583, Sakura) by freezing on dry ice. Blocks were sectioned at 7 μm thickness and longitudinal sections were mounted on glass slides (12-550-015, Fisherbrand). RNA probes from Advanced Cell Diagnostics for mouse *Des* (407921-C2), *Tnnt2* (418681-C3), *Dlx5* (478151-C3), and *Msx2* (421851-C2) were applied, in situ hybridization was performed, and the probes labeled with TSA Vivid dye Fluorophore 570 (for *Des* and *Msx2*) or 650 (for *Tnnt2* and *Dlx5*) using the RNAscope Multiplex Fluorescent Reagent Kit v2 (323270, Advanced Cell Diagnostics) according to the manufacturer's instructions. Images were acquired with a confocal microscope (SP8, Leica).

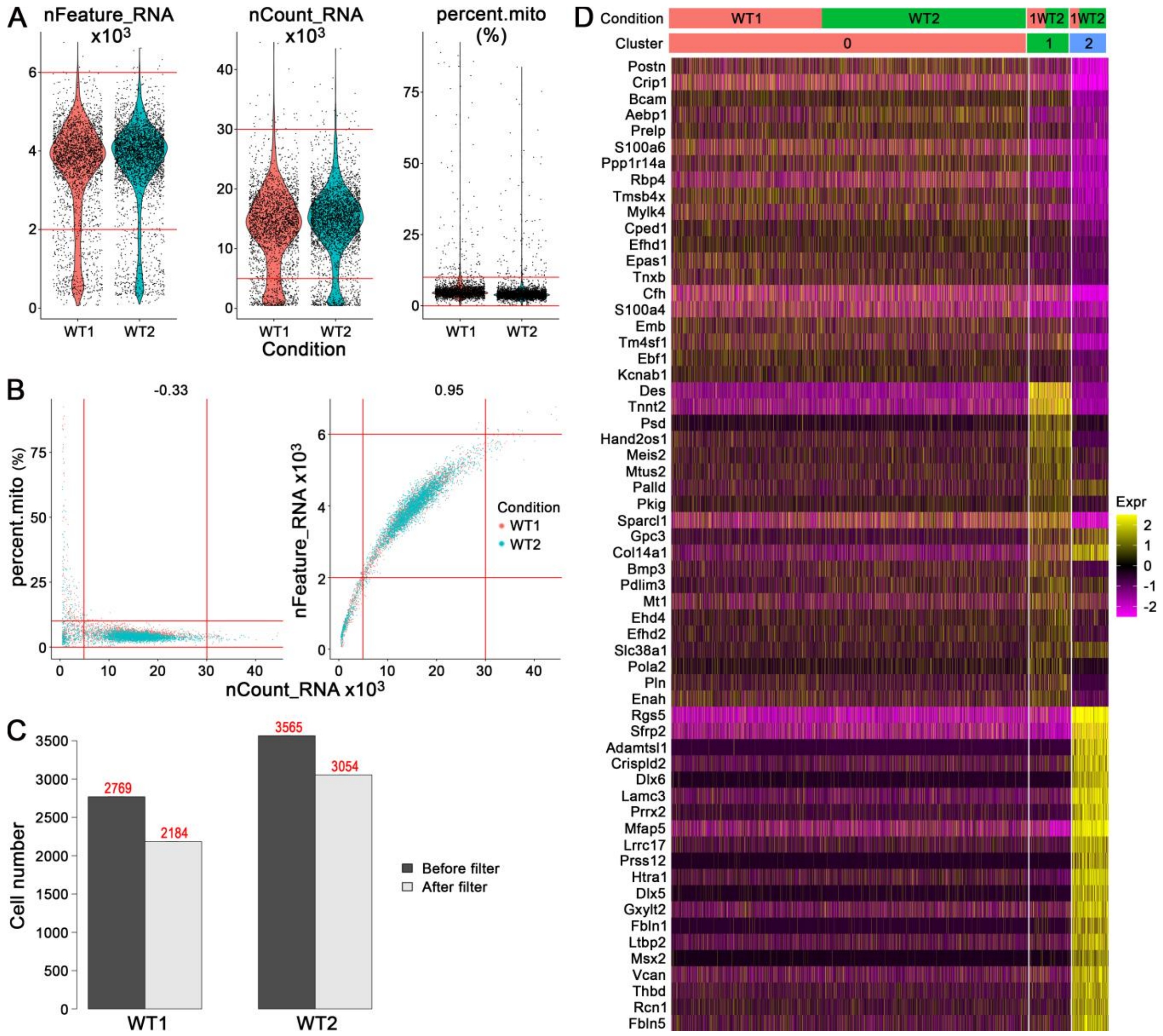


**Supplemental Figure 1: scRNA-seq of Myh11 lineage-marked SMCs of thoracic aorta.** GFP+ SMCs from root, ascending, and arch aortic segments of tamoxifen-treated, 12-week-old Myh11-CreER;mT/mG mice (referred to as WT) were analyzed by scRNA-seq. (**A**) Quality control parameters of number of genes detected per cell (nFeature_RNA), total number of unique molecular identifiers detected per cell (nCount_RNA), and fraction of counts from mitochondrial genes per cell (percent.mito), and (**B**) the parameters as covariates with Pearson correlations; threshold limits marked by red lines. (**C**) Number of cells before and after filtering based on quality control metrics. (**D**) Relative expression (expr) of top 20 markers by cluster (#0-2) and condition (2 replicates).

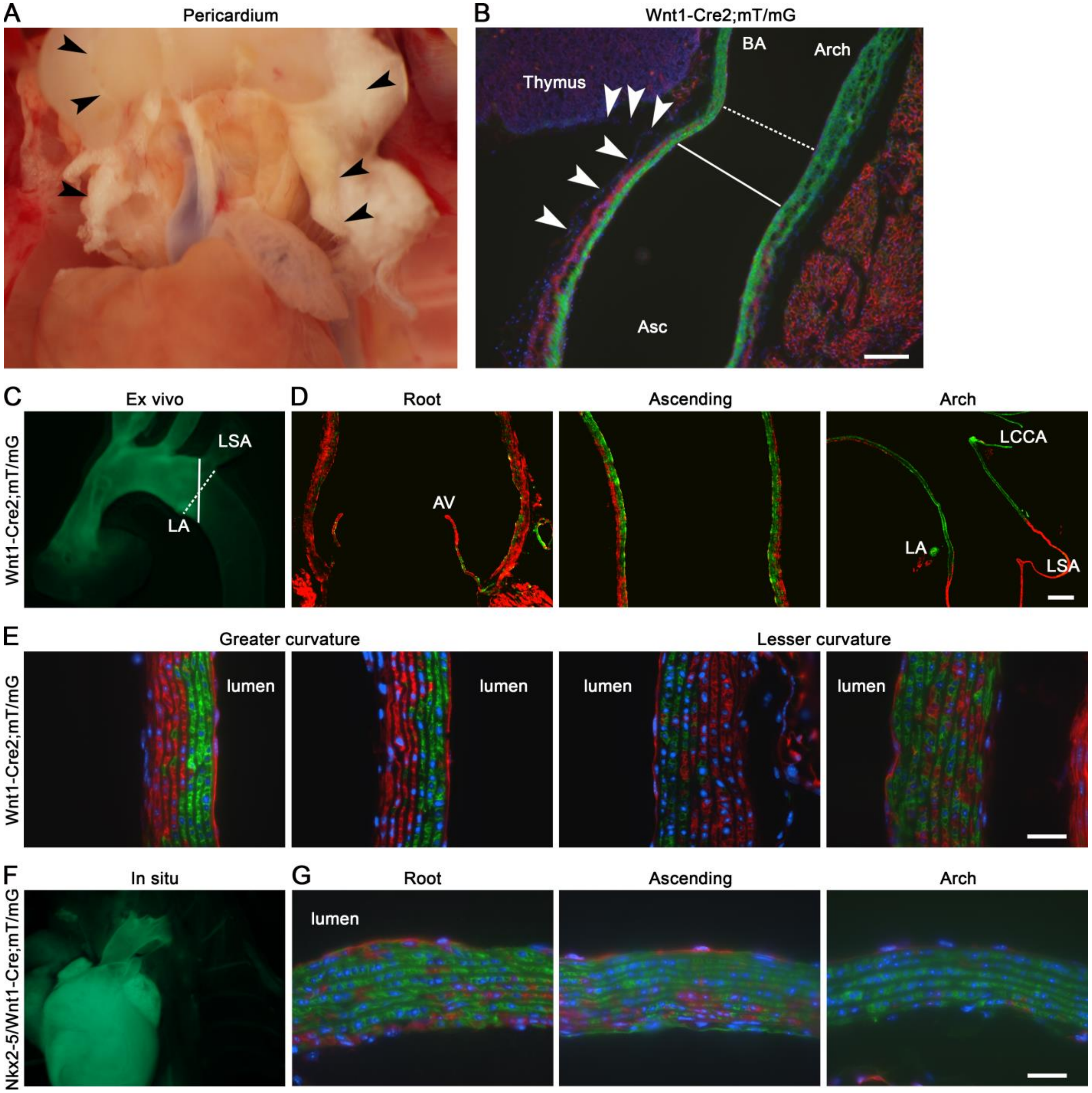


**Supplemental Figure 2: Distribution of cardiac- and neural crest-derived SMCs.** The thoracic aorta was analyzed by stereo and fluorescence microscopy in 3-week-old reporter mice. (**A**) Pericardium (arrows) after thoracotomy and prior to dissecting aorta from surrounding structures. (**B**) GFP, RFP, and DAPI fluorescence (green, red, and blue, respectively) in longitudinal aortic section of Wnt1-Cre2;mT/mG mice showing reflection of pericardium (arrows) from ascending (Asc) aorta onto thymus that corresponds with the distal extent of RFP-expressing, cardiac-derived SMCs (solid line) prior to the anatomical boundary (dashed line) of ascending and arch segments before brachiocephalic artery (BA); scale bar = 100 µm. (**C**) Thoracic aorta from Wnt1-Cre2;mT/mG mice ex vivo showing distal extent of GFP-expressing neural crest-derived SMCs (solid line) before left subclavian artery (LSA) on outer curvature and after ligamentum arteriosum (LA) on inner curvature that differs from the anatomical boundary (dashed line) of arch and descending segments after left subclavian artery; AV: aortic valve, LCCA: left common carotid artery. (**D**) Longitudinal sections of root, ascending, and arch segments from Wnt1-Cre2;mT/mG mice showing only scattered GFP-expressing neural crest-derived SMCs in the aortic root compared to around half the medial cells of the ascending aorta and all the medial cells of the aortic arch; scale bar = 100 µm. (**E**) Longitudinal sections of ascending aorta from Wnt1-Cre2;mT/mG mice showing irregular distribution of cardiac- and neural crest-derived SMCs in parts of the lesser curvature compared to typical inner neural crest-derived SMCs and outer cardiac-derived SMCs in the greater curvature; vertical orientation with lumen delineated, scale bar = 25 µm. (**F**) Heart and thoracic aorta in situ and (**G**) longitudinal aortic sections of compound Nkx2-5-Cre;Wnt1-Cre2;mT/mG mice showing absent GFP in descending thoracic aorta but occasional RFP+ medial cells of root, ascending, and arch segments; horizontal orientation with intima above, scale bar = 25 µm.

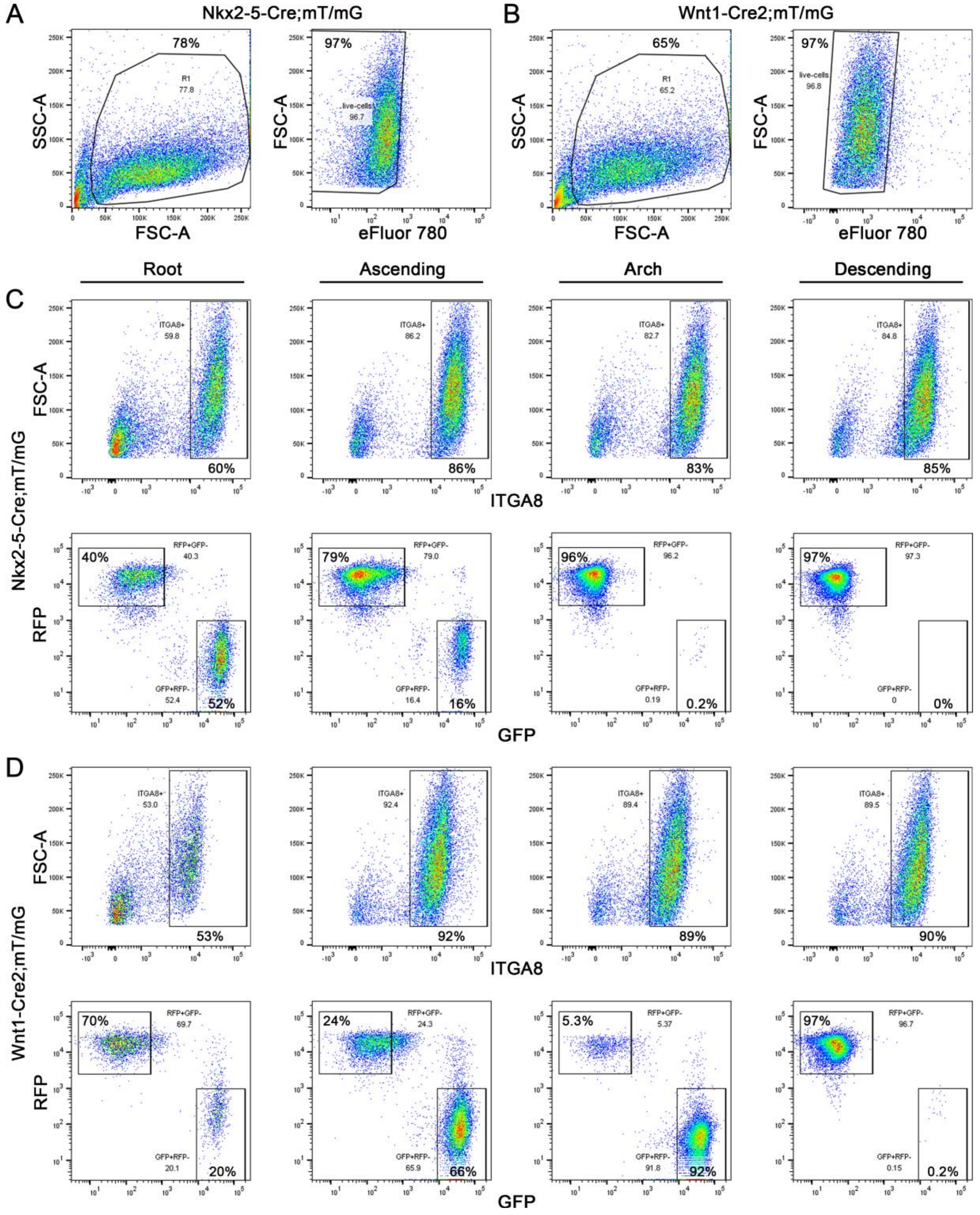


**Supplemental Figure 3: FACS sorting of cells isolated from aortic segments of cardiac and neural crest reporter mice.** Vessel wall cells were enzymatically isolated from root, ascending, arch, and descending aortic segments of 12-week-old reporter mice. Viable cells were selected by forward (FSC-A) and side (SSC-A) scatter area parameters and exclusion of eFluor 780 dye in (**A**) Nkx2-5-Cre;mT/mG and (**B**) Wnt1-Cre2;mT/mG strains. For each aortic segment, SMCs were selected by expression of integrin α8 (ITGA8) and either Cre-induced GFP expression or basal RFP expression in (**C**) Nkx2-5-Cre;mT/mG and (**D**) Wnt1-Cre2;mT/mG strains. The percent of selected cell populations of total events is shown; double positive or double negative GFP and RFP cells were not analyzed. Root cells of Nkx2-5-Cre;mT/mG mice and ascending cells of Nkx2-5-Cre;mT/mG and Wnt1-Cre2;mT/mG mice were submitted for scRNA-seq as separate GFP+ and RFP+ populations. Root cells of Wnt1-Cre2;mT/mG mice were submitted for scRNA-seq as a combined GFP/RFP+ population because of insufficient numbers of neural crest-derived SMCs. GFP+ cells from arch and descending segments of Nkx2-5-Cre;mT/mG mice and RFP+ cells from arch segments of Wnt1-Cre2;mT/mG mice were discarded. RFP+ cells from descending segments of Wnt1-Cre2;mT/mG mice were not included in scRNA-seq analysis to not unduly impact SMC clustering with cells not of cardiac nor neural crest origin.

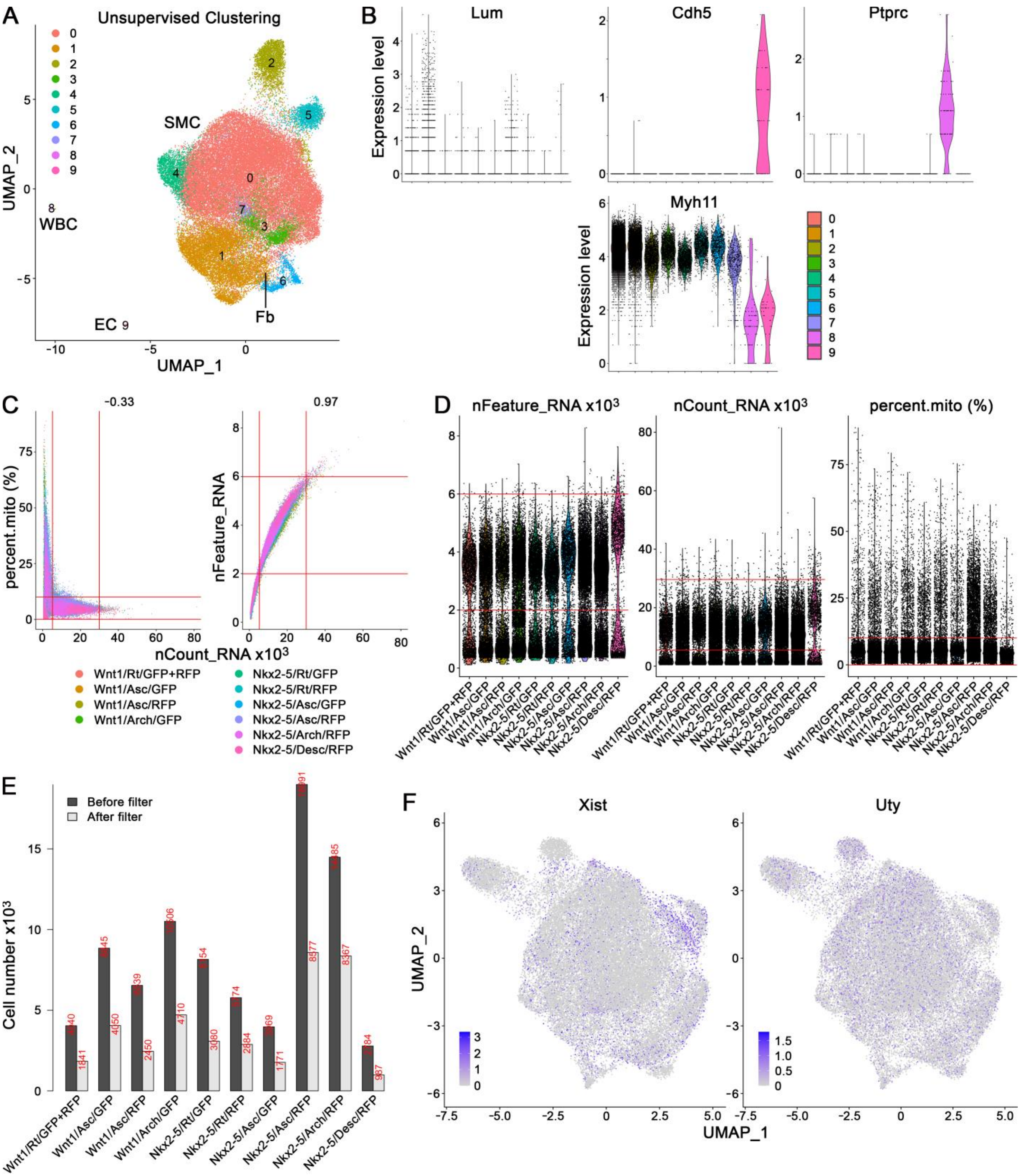


**Supplemental Figure 4: scRNA-seq of aortic SMCs.** Integrin α8+ SMCs from root (Rt), ascending (Asc), arch, and descending (Desc) aortic segments of 12-week-old Nkx2-5-Cre;mT/mG (Nkx2-5) and Wnt1-Cre2;mT/mG (Wnt1) mice were analyzed by scRNA-seq. (**A**) Clustering of all FACS-selected cells (similar to that of enriched SMCs in Figure 3A with common, cardiac subset, and neural crest subset clusters) reveals small number of fibroblasts (Fb) in part of Cluster 1, endothelial cells (EC) in Cluster 8, and leukocytes (WBC) in Cluster 9 identified by (**B**) cell type markers *Lum*, *Cdh5*, and *Ptprc* with corresponding low/absent SMC marker, *Myh11*. Therefore, cells expressing fibroblasts markers, *Lum* and/or *Dcn*, endothelial markers, *Cdh5* and/or *Pecam1*, and leukocyte markers, *Ptprc* and/or *Fcer1g* were excluded to computationally enrich SMCs. (**C**) Quality control parameters for enriched SMCs of number of genes detected per cell (nFeature_RNA), total number of unique molecular identifiers detected per cell (nCount_RNA), and fraction of counts from mitochondrial genes per cell (percent.mito) as covariates with Pearson correlations, and (**D**) as univariate parameters; threshold limits marked by red lines. (**E**) Number of cells before and after filtering by quality control metrics. (**F**) Feature plots for X chromosome gene, *Xist* (expressed preferentially in females) and Y chromosome gene, *Uty* (expressed in males) with skewed and uniform distribution, respectively.

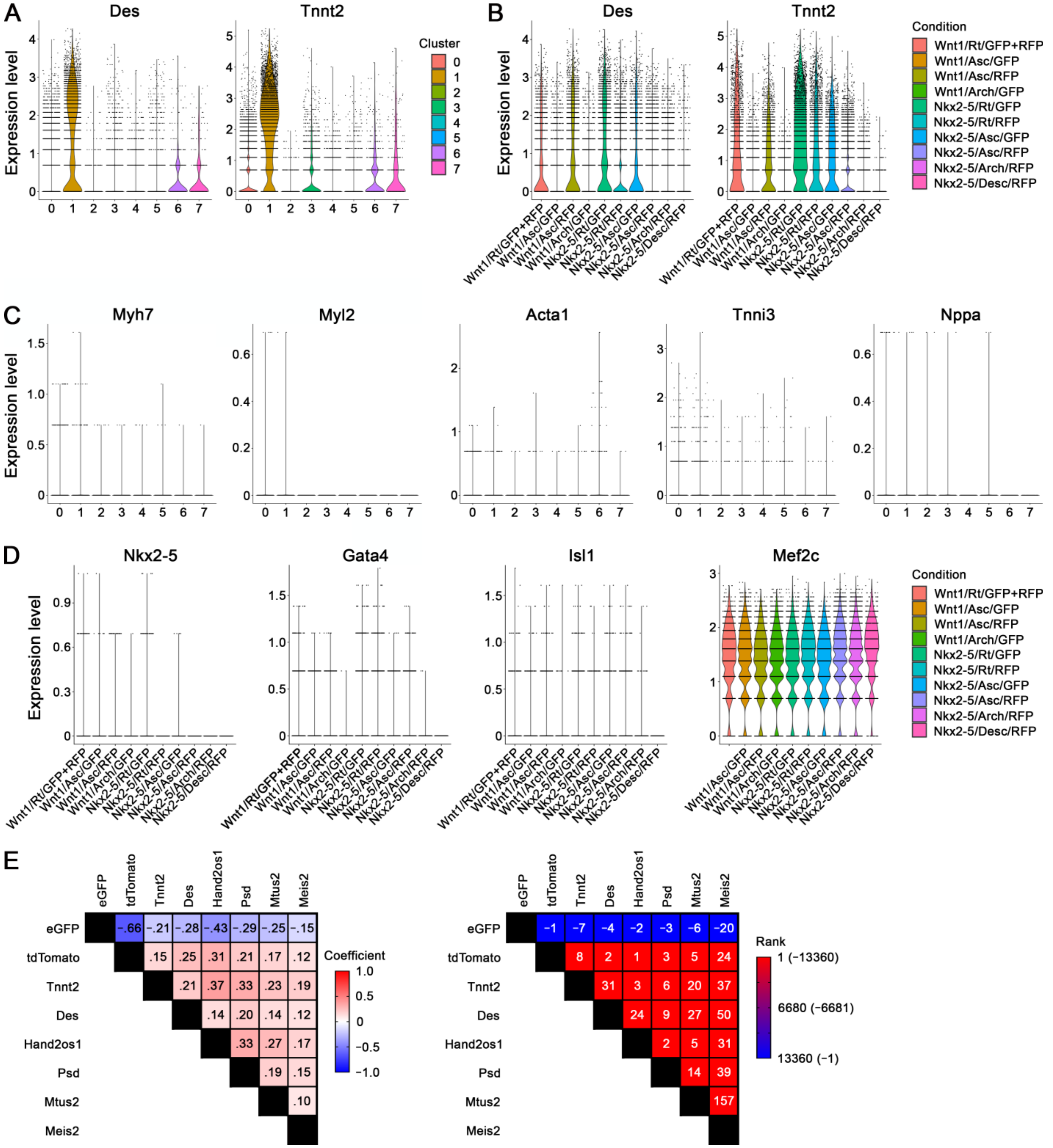


**Supplemental Figure 5: Differential gene expression by cardiac-derived SMCs.** GFP+ and RFP+ SMCs of root (Rt), ascending (Asc), arch, and descending (Desc) aortic segments from 12-week-old Nkx2-5-Cre;mT/mG (Nkx2-5) and Wnt1-Cre2;mT/mG (Wnt1) mice were analyzed by scRNA-seq analysis to determine transcript markers indicative of cardiac origin. Detection of *Des* and *Tnnt2* by (**A**) cluster and (**B**) condition revealed increased expression in cardiac subset Cluster 1 and preferential expression by cardiac-derived (i.e., Nkx2-5/GFP or Wnt1/RFP) cells in root and ascending segments. (**C**) Minimal expression of characteristic cardiomyocyte transcripts, *Myh7*, *Myl2*, *Acta1*, *Tnni3*, and *Nppa* suggest that contamination of SMC transcripts by that of other cell types is unlikely the basis for readily detectable *Des* and *Tnnt2*. (**D**) *Nkx2-5* is minimally detected, *Gata4* has decreasing expression gradient from proximal to distal aorta in both cardiac- and neural crest-derived SMCs, *Isl1* is equally detected in cardiac- and neural crest-derived, but not descending segment, SMC, and *Mef2c* is ubiquitously expressed by adult SMCs. (**E**) Gene co-expression analysis of selected transcripts in root SMCs from Wnt1-Cre2;mT/mG mice (which had not been sorted as separate GFP+ and RFP+ cell populations) demonstrated positive correlation of *tdTomato* and inverse correlation of *eGFP* to the cardiac SMC markers, *Tnnt2*, *Des*, *Hand2os1*, *Psd*, *Mtus2*, and *Meis2*; color-coded matrices for Pearson correlation coefficient and correlation rank among 13,361 total genes.

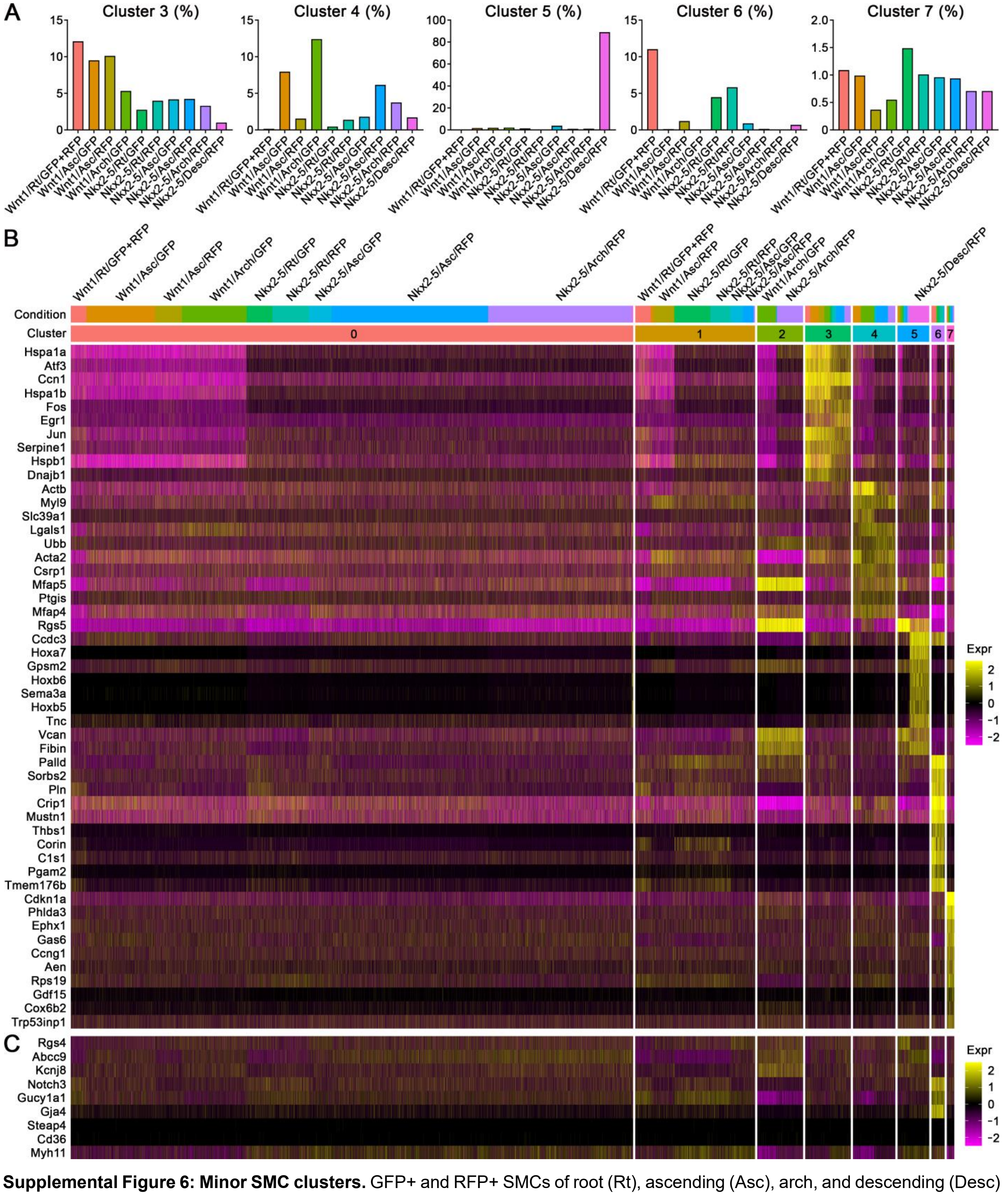


**Supplemental Figure 6: Minor SMC clusters.** GFP+ and RFP+ SMCs of root (Rt), ascending (Asc), arch, and descending (Desc) aortic segments from 12-week-old Nkx2-5-Cre;mT/mG (Nkx2-5) and Wnt1-Cre2;mT/mG (Wnt1) mice were analyzed by scRNA-seq. (**A**) Contributions of SMCs from different aortic segments to minor clusters #3-7. (**B**) Cluster 3 is distinguished by immediate early genes (including *Hspa1a*, *Atf3*, *Ccn1*, *Haspa1b*, *Fos*, and *Egr1*) indicative of SMC activation. Cluster 4 has increased expression of several cytoskeleton and contractile genes (*Actb*, *Myl9*, and *Acta2*). Cluster 5 from descending aorta is distinguished by distal Hox genes (*Hoxa7*, *Hoxb6*, and *Hoxb5*). Cluster 6 has genes preferentially expressed by microvascular SMCs (*Sorbs2*, *Crip1*, and *C1s1*). Cluster 7 has several genes associated with cell cycle/survival (*Cdkn1a*, *Phlda3*, and *Ccng1*). (**C**) Cluster 5 not from descending aorta lacked distal Hox genes but expressed certain microvascular-related markers (*Rgs4*, *Abcc9*, and *Kcnj8* in addition to *Rgs5* in panel B) but not other microvascular markers expressed in Cluster 6 (*Notch3*, *Gucy1a1*, and *Gja4* in addition to *Sorbs2*, *Crip1*, and *C1s1* in panel B). Both Cluster 5 and 6, however, did not express a complete repertoire of pericyte markers (*Steap4* and *Cd36*) but expressed the archetypal SMC marker (*Myh11*) supporting categorization as microvascular SMCs.

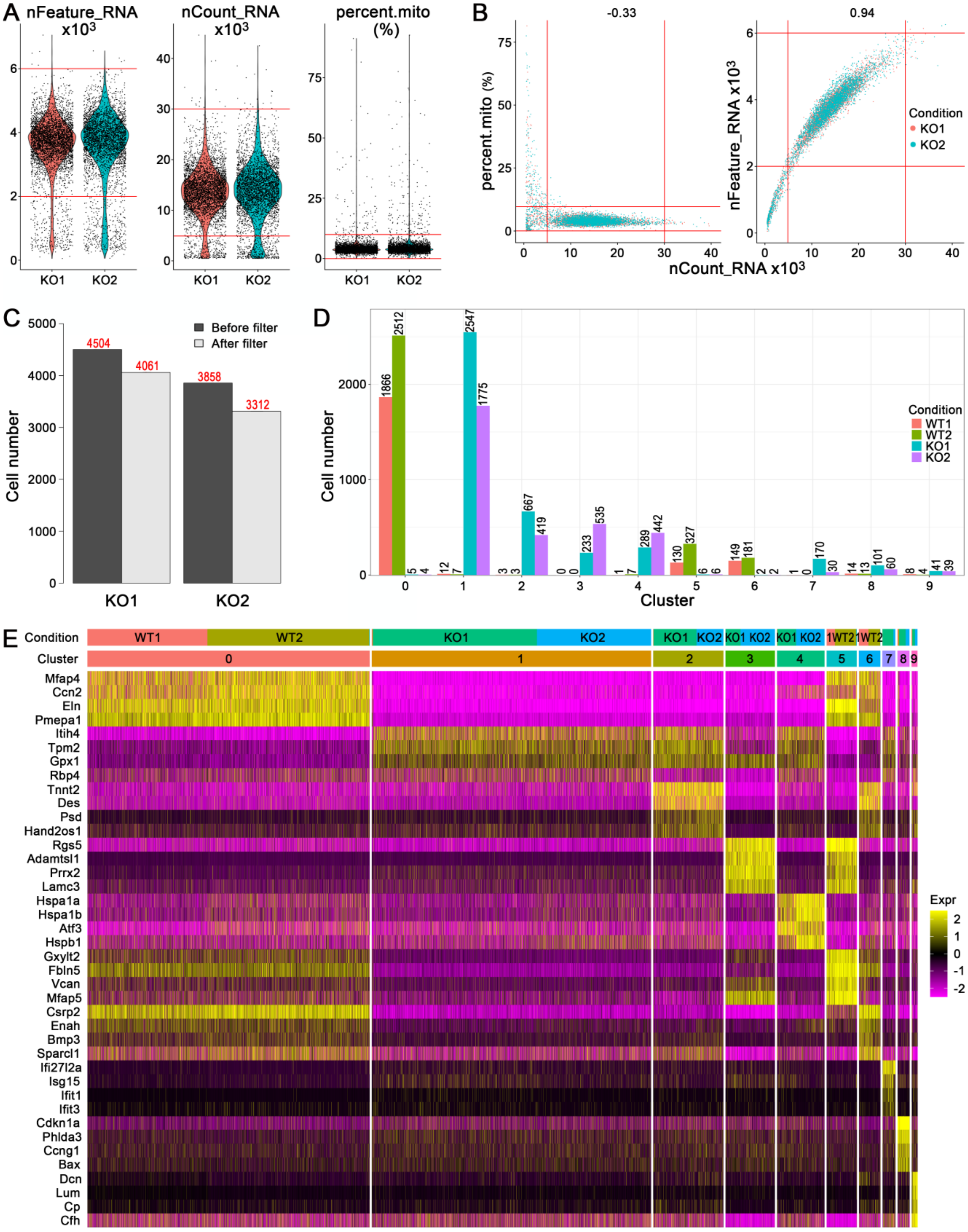


**Supplemental Figure 7: scRNA-seq of aortic SMCs after TGFβ signaling disruption.** GFP+ SMCs from root, ascending, and arch aortic segments of tamoxifen-treated, 12-week-old Myh11-CreER;mT/mG mice (WT) and Tgfbr1$^{f/f}$/Tgfbr2$^{f/f}$;Myh11-CreER;mT/mG (KO) mice were analyzed by scRNA-seq. (**A**) Number of genes detected per cell (nFeature_RNA), total number of unique molecular identifiers detected per cell (nCount_RNA), and fraction of counts from mitochondrial genes per cell (percent.mito), (**B**) parameters as covariates with Pearson correlations, threshold limits marked by red lines, and (**C**) number of cells before and after quality control filtering for KO cells (metrics for WT cells shown in Supplemental Figure 1). (**D**) Number of WT and KO cells per cluster. (**E**) Relative expression (expr) of top 4 markers by cluster (#0-9) and condition (2 WT and 2 KO).

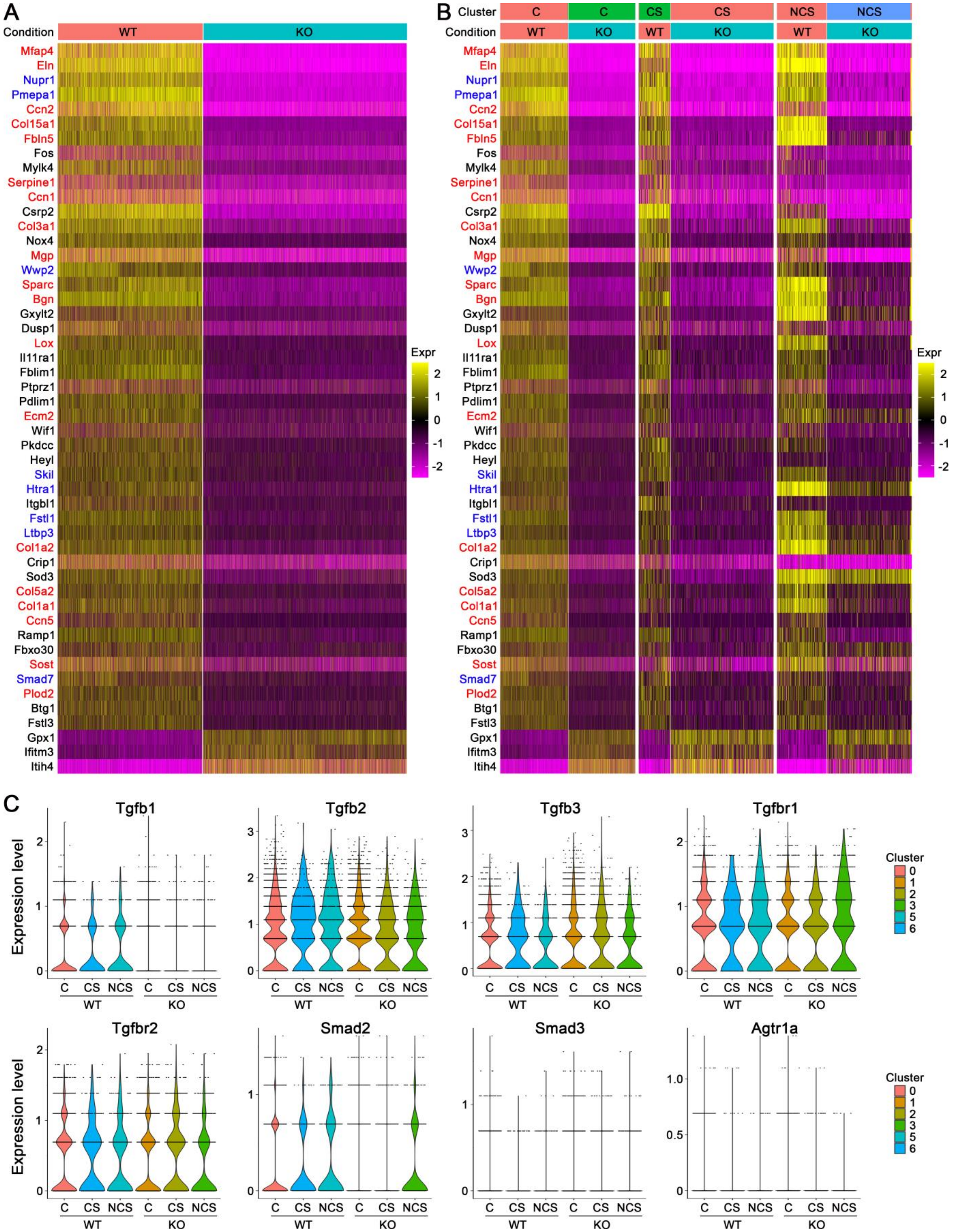


**Supplemental Figure 8: TGFβ-dependent genes in cardiac- vs. neural crest-derived SMCs.** GFP+ SMCs from root, ascending, and arch aortic segments of tamoxifen-treated, 12-week-old Myh11-CreER;mT/mG (WT) and Tgfbr1$^{f/f}$/Tgfbr2$^{f/f}$;Myh11-CreER;mT/mG (KO) mice were analyzed by scRNA-seq. (**A**) Relative expression (expr) of top 50 markers by condition (WT and KO) and (**B**) separate analyses of common (C), cardiac subset (CS), and neural crest subset (NCS) clusters; ECM genes highlighted in red and TGFβ signaling regulator genes in blue font. (**C**) Expression of TGFβ and angiotensin II signaling effectors, *Tgfb1*, *Tgfb2*, *Tgfb3*, *Tgfbr1*, *Tgfbr2*, *Smad2*, *Smad3*, and *Agtr1a*, in common, cardiac subset, and neural crest subset clusters.

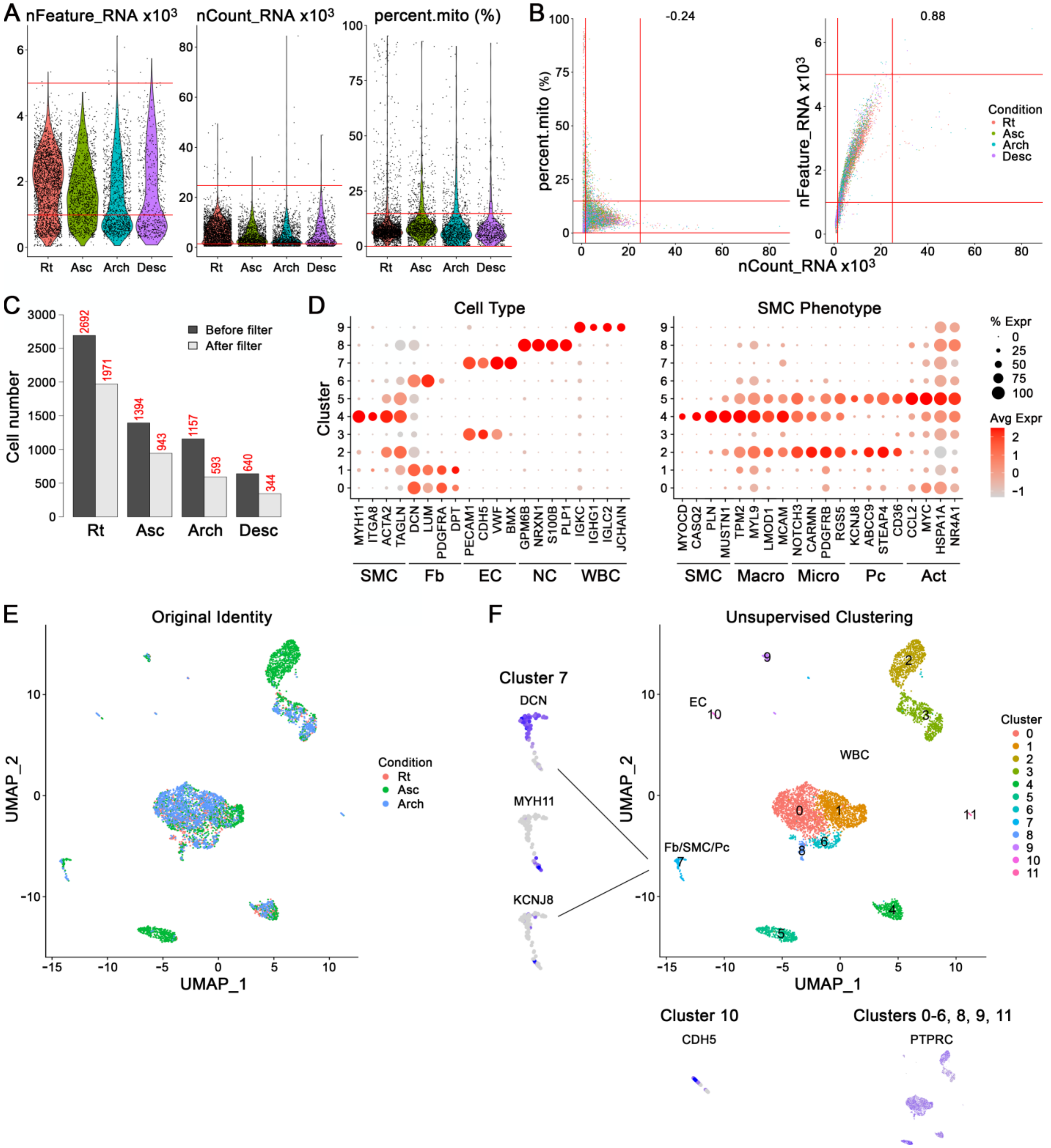


**Supplemental Figure 9: scRNA-seq of human aortic SMCs.** CD45-/CD235a- vessel wall cells from root (Rt), ascending (Asc), arch, and descending (Desc) media specimens of 55-year-old, male organ donor were analyzed by scRNA-seq. (**A**) Number of genes detected per cell (nFeature_RNA), total number of unique molecular identifiers detected per cell (nCount_RNA), and fraction of counts from mitochondrial genes per cell (percent.mito) and (**B**) parameters as covariates with Pearson correlations, threshold limits marked by red lines. (**C**) Number of cells before and after quality control filtering. (**D**) expression (expr) of cell type markers for SMCs/pericytes, fibroblasts, endothelial cells, neuronal cells, and leukocytes, and SMC phenotype markers categorized as aortic SMC-specific, predominantly macrovascular (Macro), predominantly microvascular (Micro), pericyte-specific (Pc), and activated (Act). Alternatively, vessel wall cells not depleted of leukocytes or erythrocytes from full thickness aorta specimens (including adventitia) of 51-year-old, female subject were analyzed by scRNA-seq with (**E**) original identity by aortic segment and (**F**) unsupervised clustering into 12 clusters of leukocytes (WBC; #0-6, 8, 9, 11), fibroblasts/SMCs/pericytes (Fb/Pc; #7), and endothelial cells (EC; #10) with feature plot insets for *DCN*, *MYH11*, and *KCNJ8* for fibroblasts, SMCs, and pericytes, respectively, in Cluster 7, *CDH5* for endothelial cells in Cluster 10, and *PTPRC* for leukocytes in Clusters 0-6, 8, 9, and 11.
